# Computational genomics of brain tumors: identification and characterization of glioma candidate biomarkers through multi-omics integrative molecular profiling

**DOI:** 10.1101/798785

**Authors:** Lin Liu, Guangyu Wang, Liguo Wang, Chunlei Yu, Mengwei Li, Shuhui Song, Lili Hao, Lina Ma, Zhang Zhang

**Author notes:** Corresponding authors (M.L.N.), (Z.Z.). The Methodist Hospital Research Institute, 6670 Bertner Ave, Houston, Texas 77030, United States. These authors contributed equally.

## Abstract

Glioma is one of the most common malignant brain tumors and exhibits low resection rate and high recurrence risk. Although a large number of glioma studies powered by high-throughput sequencing technologies have led to massive multi-omics datasets, there lacks of comprehensive integration of glioma datasets for uncovering candidate biomarker genes. In this study, we collected a large-scale assemble of multi-omics multi-cohort datasets from worldwide public resources, involving a total of 16,939 samples across 19 independent studies. Through comprehensive multi-omics molecular profiling across different datasets, we revealed that *PRKCG* (Protein Kinase C Gamma), a brain-specific gene detectable in cerebrospinal fluid, is closely associated with glioma. Specifically, it presents lower expression and higher methylation in glioma samples compared with normal samples. *PRKCG* expression/methylation change from high to low is indicative of glioma progression from low-grade to high-grade and high RNA expression is suggestive of good survival. Importantly, *PRKCG* in combination with *MGMT* is effective to predict survival outcomes after TMZ chemotherapy in a more precise manner. Collectively, *PRKCG* bears the great potential for glioma diagnosis, prognosis and therapy, and *PRKCG*-like genes may represent a set of important genes associated with different molecular mechanisms in glioma tumorigenesis. Accordingly, our study indicates the importance of computational integrative multi-omics data analysis and represents a data-driven scheme toward precision tumor subtyping and accurate personalized healthcare.

**Author Summary:** Glioma is a type of brain tumors that represents one of the most lethal human malignancies with little chance for recovery. Nowadays, more and more studies have adopted high-throughput sequencing technologies to decode the molecular profiles of glioma from different omics levels, accordingly resulting in massive glioma datasets generated from different projects and laboratories throughout the world. Therefore, it has become crucially important on how to make full use of these valuable datasets for computational identification of glioma candidate biomarker genes in aid of precision tumor subtyping and accurate personalized treatment. In this study, we comprehensively integrated glioma datasets from all over the world and performed multi-omics molecular data mining. We revealed that *PRKCG*, a brain-specific gene abundantly expressed in cerebrospinal fluid, bears the great potential for glioma diagnosis, prognosis and treatment prediction, which has been consistently observed on multiple independent datasets. In the era of big data, our study highlights the significance of computational integrative data mining toward precision medicine in cancer research.

## Introduction

Glioma, one of the serious central nervous system (CNS) tumors, represents ∼80% of malignant brain tumors (1, 2) and exhibits low resection rate and high recurrence risk (3). Since tumor classification benefits accurate diagnosis and facilitates precise treatment, gliomas can be classified, according to the histologic grading schemes, into LGG (astrocytoma, oligodendroglioma and oligoastrocytoma) and GBM (glioblastoma multiforme) (4). Therefore, identification of reliable molecular biomarkers for precise classification of different-grade gliomas is crucial to aid tumor diagnosis, establish appropriate therapies, recognize prognostic outcome and predict therapeutic response (5).

Powered by high-throughput sequencing technologies, a set of molecular biomarkers have been discovered from different omics levels to assist glioma diagnosis and treatment (6, 7). Among them, isocitrate dehydrogenase (*IDH*) mutation and 1p/19q co-deletion (codel) are two most important genetic events for glioma grading (8–10). Patients with *IDH* mutation (*IDH*-mut) have longer survival than those with *IDH* wild-type (*IDH*-WT) (11, 12). And the 1p/19q codel is a distinctive feature of oligodendroglioma (13, 14). Furthermore, based on these two genetic alterations, accumulated evidence suggested that gliomas can be divided into three subtypes (*IDH*-mut & 1p/19q codel, *IDH*-mut & 1p/19q non-codel, and *IDH*-WT & 1p/19q non-codel), which are associated with diverse clinical outcomes (15). Accordingly, in 2016, the World Health Organization (WHO), in light of both histology and significant genetic events (mainly by *IDH* and 1p/19q), divided gliomas into five categories (16, 17), including three LGGs (diffuse astrocytoma, *IDH*-mut & 1p/19q non-codel; oligodendroglioma, *IDH*-mut & 1p/19q codel; diffuse astrocytoma, *IDH*-WT & 1p/19q non-codel) and two GBMs (*IDH*-mut; *IDH*-WT). Meanwhile, molecular markers at the transcriptome level have also been identified (18–21); for example, an overexpression of epidermal growth factor receptor variant III (*EGFRvIII*) has been reported to associate with malignant progression of GBM (22–24). In addition, epigenetic modifications are also implicated in glioma (25–29). One classical biomarker is O6-methylguanine-DNA-methyltransferase (*MGMT*) (30); patients with methylated *MGMT* promoter have better clinical outcomes and are more sensitive to the alkylating chemotherapy than those without methylated *MGMT* promoter (31–33).

Nowadays, there is an increasing number of high-throughput studies for better understanding of glioma tumorigenesis (34–38), resulting in massive multi-omics datasets generated from different projects and laboratories throughout the world. However, there lacks of comprehensive integration of glioma datasets for computationally identifying and characterizing candidate biomarkers. To this end, we collected a large-scale assemble of multi-omics multi-cohort datasets from worldwide public resources and detected candidate biomarker genes through comprehensive integrative molecular profiling on multiple independent datasets. We revealed that *PRKCG*, a gene specifically expressed in brain and detectable in cerebrospinal fluid (CSF), is closely associated with glioma, indicative of a potential biomarker for glioma diagnosis, prognosis and treatment prediction.

## Materials and Methods

### Data collection

In this study, we collected a comprehensive assemble of multi-omics datasets (including genomics, transcriptomics, DNA methylomics and proteomics) from The Cancer Genome Atlas (TCGA, https://portal.gdc.cancer.gov/) (35), Genotype-Tissue Expression Portal (GTEx, https://gtexportal.org/home/) (39), Gene Expression Omnibus (GEO, https://www.ncbi.nlm.nih.gov/geo), Ivy Glioblastoma Atlas Project (Ivy GAP, http://glioblastoma.alleninstitute.org) (40) and Chinese Glioma Genome Atlas (CGGA, http://www.cgga.org.cn) (41, 42). Particularly, discovery datasets were derived from TCGA, GTEx and large cohort studies in GEO (GSE83710, GSE16011 and GSE36278 for protein, expression and methylation, respectively). As a result, a total of five discovery datasets and fourteen validation datasets were obtained. For convenience, each dataset collected in this study is assigned a unique accession number with the format: [D/V][*i*]-[TCGA/GTEx/GEO/CGGA/Ivy GAP]-[E/V/P/M], where D/V in the first bracket represents the dataset for discovery or validation, *i* in the second bracket indicates the dataset number, the third bracket shows the data source (as mentioned above), and the last bracket indicates the data type, namely, E for RNA expression, V for CNV, P for protein expression and M for DNA methylation, respectively. The detailed information about all collected datasets was tabulated in Table 1.

**Table 1.**
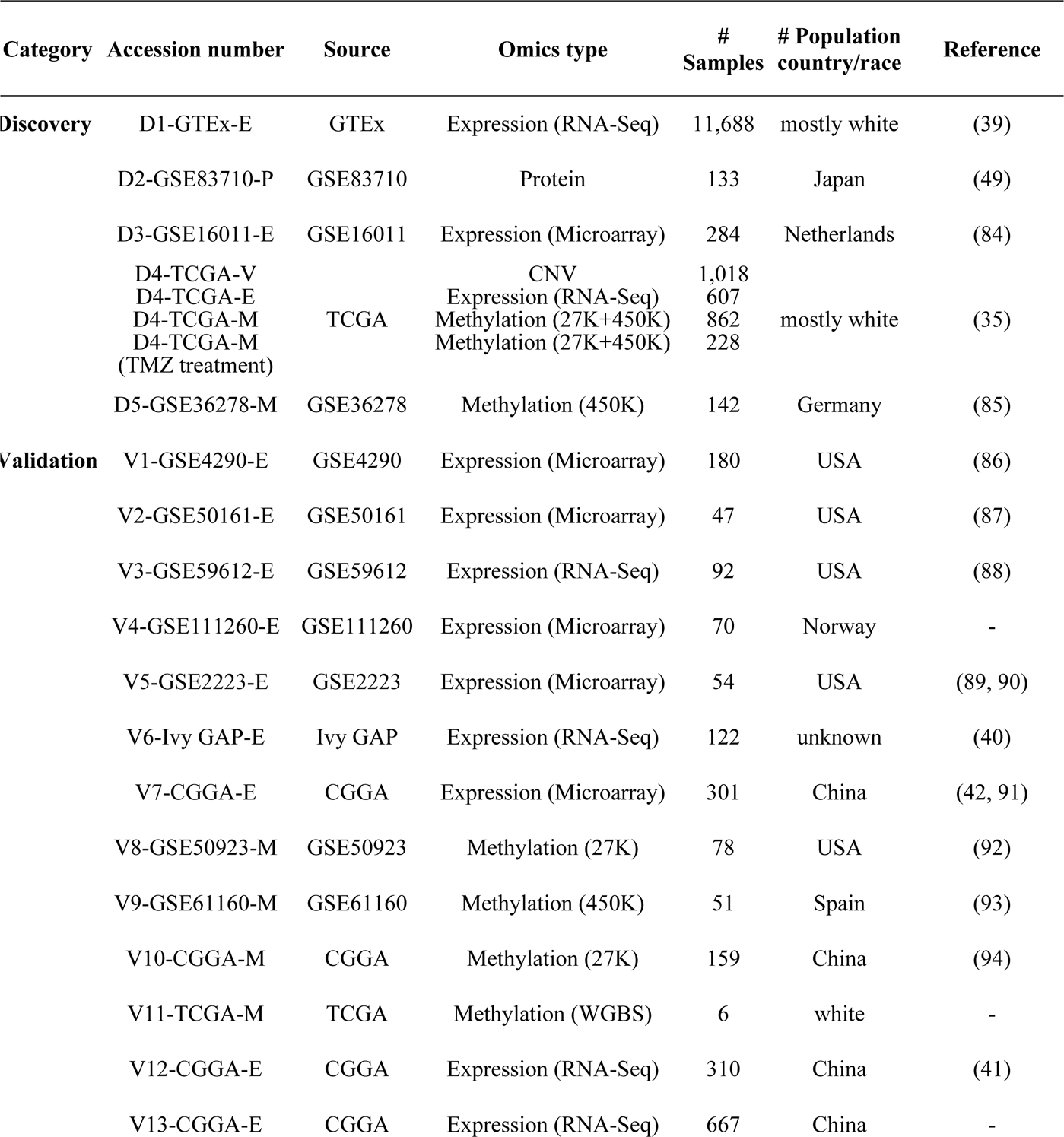
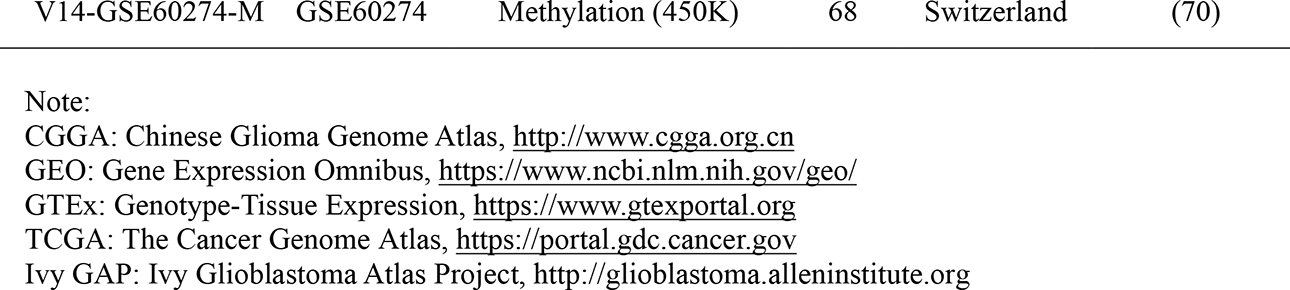
Summary of multi-omics multi-cohort glioma datasets

### Identification of brain-specific genes

To identify brain-specific genes, we used the RNA-Seq dataset from GTEx (2016-01-15; v7) (39), which contains 11,688 samples across 53 tissue sites of 714 donors. Considering that several tissues have multiple different sites, gene expression levels were averaged over sites that are from the same tissue. To reduce background noise, genes with maximal expression levels smaller than 10 TPM (Transcripts Per Million) were removed. Finally, we obtained a total of 15,176 gene expression profiles across 30 tissues (S1 Table).

Based on the expression levels across 30 tissues, we calculated the tissue specificity index τ (43) for each gene to identify tissue-specific genes. τ is valued between 0 and 1, where 0 represents housekeeping genes that are consistently expressed in different tissues, and 1 indicates tissue-specific genes that are exclusively expressed in only one tissue (43). In this study, brain-specific genes were defined as those genes that are maximally expressed in the brain with τ>0.9. As a consequence, a list of the top 100 brain-specific genes ranked by the τ index were obtained for further analysis (S2 Table).

### Sample classification

To comprehensively study the potential of *PRKCG* in glioma diagnosis, we compared the molecular profiles between normal and glioma samples, between LGG and GBM samples, between primary GBM (pGBM) and recurrent GBM (rGBM) samples, and between glioma samples with different anatomic features. We collected 122 GBM samples from the Ivy GAP database (40) and grouped them according to their anatomic regions, namely, leading edge (LE, the ratio of tumor/normal cells is about 1–3/100), infiltrating tumor (IT, the ratio of tumor/normal cells is about 10–20/100), cellular tumor (CT, the ratio of tumor/normal cells is about 100/1 to 500/1), pseudo-palisading cells around necrosis (PAN, the narrow boundary of cells along the dead tissue), and microvascular proliferation (MVP, two or more blood vessels sharing a common vessel wall of endothelial).

We investigated the prognostic role of *PRKCG* by dividing samples into subgroups based on *PRKCG*’s expression level within all glioma samples and also within LGG and GBM samples, respectively. When exploring the predictive role of *PRKCG*, we obtained DNA methylation status (methylated and unmethylated) directly from the original study (35), which was defined based on the beta value cutoff 0.3.

### Identification of *PRKCG*-like genes

Genes that satisfy the following criteria were regarded as *PRKCG*-like genes: (1) Higher methylation level of at least one CpG site (promoter region) in glioma samples than normal samples; (2) Higher DNA methylation level in LGG samples than GBM samples; (3) Higher expression level in LGG samples than GBM samples; and (4) Lower expression level in glioma samples than normal samples. As a result, we obtained a total of 542 *PRKCG*-like genes, which were further divided into two groups according to their correlations between gene expression and methylation, namely, 114 genes with negative correlation and 297 genes with positive correlation.

### Statistical analysis

All statistical analyses were performed using R version 3.3.2. The Wilcoxon test was used for the analysis of the difference in gene expression/methylation between tumor and normal samples, and between different glioma subtypes. The statistical significance levels were coded by *ns* (not significant) *p* > 0.05, * *p* < 0.05, ** *p* < 0.01 and *** *p* < 0.001. We performed the survival analysis using the Kaplan-Meier method and estimated the statistical difference using the log-rank test.

### Data availability statement

All datasets integrated in this study were obtained from multiple public database resources (see details in Table 1), which are freely available at ftp://download.big.ac.cn/glioma_data/.

## Results and Discussion

### *PRKCG* is a brain-specific gene and detectable in cerebrospinal fluid

Tissue-specific genes are believed to be crucial for identifying potential biomarkers with high specificity (44–48). To identify candidate genes with brain specificity, we integrated expression data from GTEx (D1-GTEx-E) (39), explored all genes’ expression profiles and their tissue specificity, and identified a list of top 100 brain-specific genes (S2 Table). To achieve the detectability in the periphery, we assembled a total of 1,126 CSF-detectable proteins from GEO (D2-GSE83710-P) (49), due to the critical significance of CSF as a feasible means to detect genes expressed in human CNS (50, 51). After integrating brain-specific genes with CSF proteins, we revealed that there are five brain-specific proteins that can be detected in CSF (Fig 1 and S1 Fig), in terms of fluorescence intensity from high to low, namely, *PRKCG, BCAN*, *OPCML*, *GFAP* and *CAMK2A*, which are diversely expressed in different brain regions (S2 Fig). Specifically, *BCAN*, a member of the lectican family of chondroitin sulfate proteoglycans, is highly expressed in glioma and may promote cell motility of brain tumor cells (52, 53). In addition, the fusion event between *BCAN* and *NTRK1* (*BCAN*-*NTRK1*) is a potential glioma driver and therapeutic target (54). *OPCML* encodes a member of the IgLON subfamily in the immunoglobulin proteins and is down-regulated in gliomas and other brain tumors (55, 56). *GFAP*, encoding one of the major intermediate filament proteins of mature astrocytes (57), can be used to assess the differentiation state of astrocytoma (58). *CAMK2A* is a calcium calmodulin-dependent protein kinase and reduced expression of *CAMK2A* is associated with better survival in GBM (59, 60).

**Fig 1.**
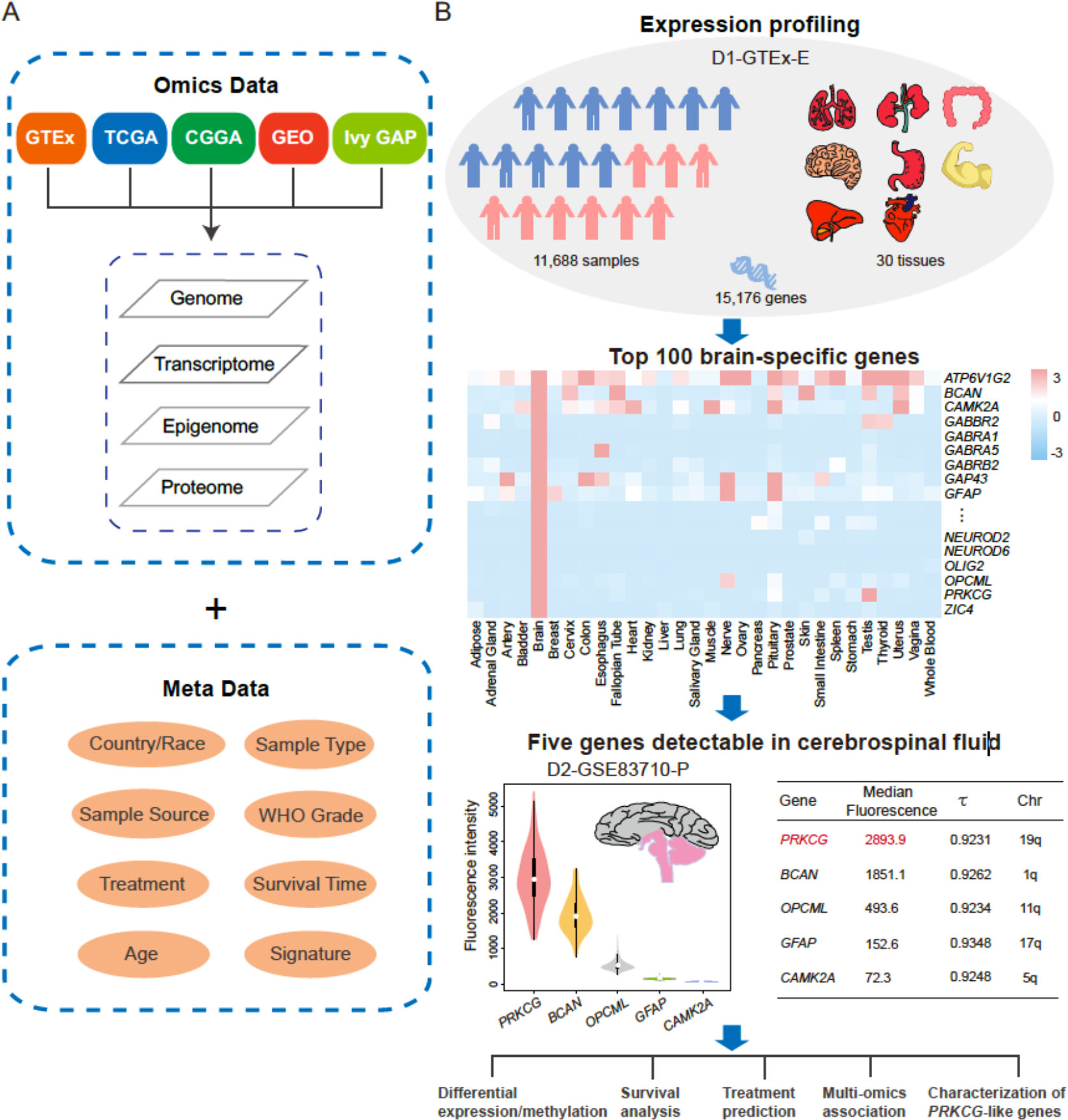
Datasets and bioinformatic analysis workflow. (A) A comprehensive assemble of multi-omics datasets and their corresponding meta data were integrated from GTEx, TCGA, CGGA, GEO and Ivy GAP. **(B)** An integrative analysis workflow was adopted, including detection of brain-specific genes, identification of CSF-detectable genes, ranking of candidate genes in light of protein fluorescence. A series of bioinformatic analyses were performed, including differential expression/methylation analysis, survival analysis, treatment prediction, multi-omics association and characterization of *PRKCG*-like genes.

Remarkably, *PRKCG* (Protein Kinase C Gamma), a member of protein kinase C (PKC) family located in 19q, exhibits higher fluorescence intensity than the other four genes (Fig 1 and S1 Fig). The expression profile of *PRKCG* across multiple brain developmental stages reveals that its expression is extremely lower in the prenatal stages, but dramatically increases in the infancy stages and is stabilized in the latter stages according to GenTree (S3 Fig) (61). Previous studies have documented that unlike other PKC family members that are expressed in many tissues aside from brain, *PRKCG* is brain-specifically expressed (62) and that mutations in *PRKCG* are associated with spinocerebellar ataxia (63, 64). Additionally, it has been reported that PKC signaling pathways contribute to the aggressive behavior of glioma cells (65) and atypical PKC isozymes are fundamental regulators of tumorigenesis (66). To our knowledge, several genes in 19q are closely associated with glioma (e.g., *TTYH1*, *UBE2S* (67, 68)). However, the potential role of *PRKCG* in glioma remains unknown, and therefore, comprehensive molecular characterization of *PRKCG* across multi-omics glioma datasets is highly desirable.

### *PRKCG* is significantly differentially expressed among normal, LGG and GBM samples

We first investigated the expression pattern of *PRKCG* among normal, LGG and GBM samples by using multiple discovery and validation datasets. We found that *PRKCG* expression is significantly reduced in gliomas by contrast to normal samples (Fig 2A-F; *p-*value < 0.01, Wilcoxon test). Furthermore, we discovered that *PRKCG* shows significantly different expression profiles among different anatomic regions (Fig. 2G; *p-*value < 0.01, Wilcoxon test). Strikingly, *PRKCG* expression is highest in LE (the outermost boundary of the tumor), decreased in IT (the intermediate zone between the LE and the serious CT regions), and lowest in the serious regions (CT, PAN and MVP) (see details in Materials and Methods). Consistently, comparison between different-grade gliomas showed that *PRKCG* expression is significantly lower in GBM samples than LGG samples (Fig 2H-J; *p-*value < 0.01, Wilcoxon test). We further investigated its expression across pan-cancer samples. Although it has been documented that *PRKCG* is up-regulated in colon cancer (69), the up-regulation in colon cancer is extremely lower by comparison with glioma (LGG and GBM) (S4 Fig). Taken together, these results presumably suggest that *PRKCG* is closely associated with glioma and its reduced expression is coupled with glioma progression (Fig 2), highlighting its possible potential for glioma diagnosis.

**Fig 2.**
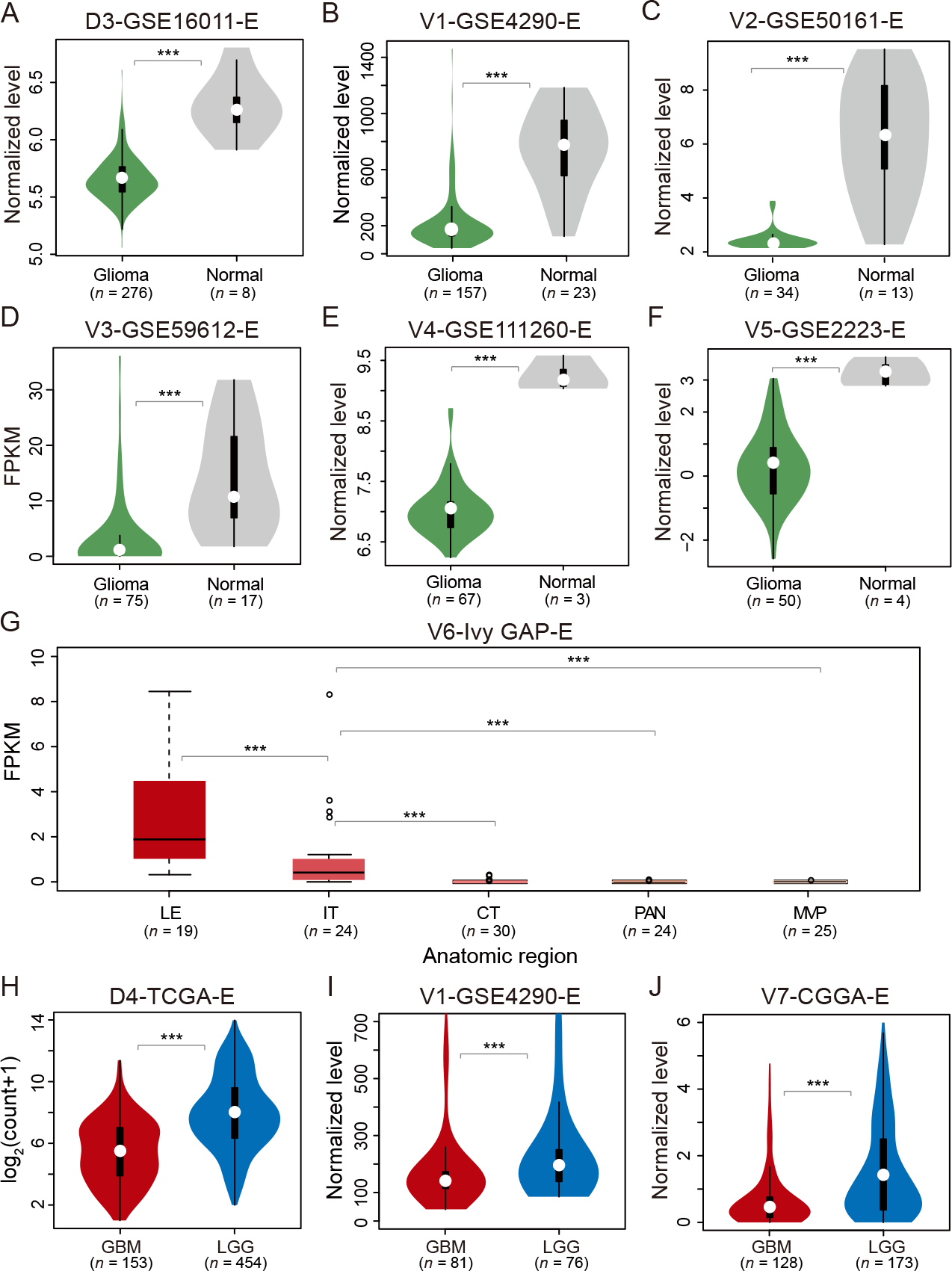
Expression profiles of *PRKCG* across normal, LGG and GBM samples. *PRKCG* expression profiles were compared between glioma and normal samples (D3-GSE16011-E in panel A [RMA normalized], V1-GSE4290-E in panel B [MAS5 normalized], V2-GSE50161-E in panel C [gcRMA normalized], V3-GSE59612-E in panel D, V4-GSE111260-E in panel E [RMA normalized], V5-GSE2223-E in panel F [Lowess normalized]), between different anatomic regions (V6-Ivy GAP-E in panel G), and between GBM and LGG samples (D4-TCGA-E in panel H, V1-GSE4290-E in panel I [MAS5 normalized] and V7-CGGA-E in panel J [Lowess normalized]). All the normalization methods labeled above were derived from and detailed in their corresponding publications, and all these datasets were made publicly accessible at ftp://download.big.ac.cn/glioma_data/. The Wilcoxon tests were performed and the statistical significance levels were coded by: *ns p*>0.05, * *p*<0.05, ** *p*<0.01 and *** *p*<0.001.

### *PRKCG* expression is highly sensitive to survival

*PRKCG* expression change from high to low is indicative of progression from normal to glioma and from LGG to GBM (Fig 2), implying that *PRKCG* expression is significantly associated with glioma progression. Importantly, we observed that *PRKCG* expression is significantly associated with survival rate, which is testified by multiple independent datasets (Fig 3). Specifically, higher expression of *PRKCG* is indicative of longer overall survival in all glioma samples (Fig 3A-B; *p*-value < 0.01, log-rank test). When separating LGG samples from GBM samples, it is consistently observed that higher expression, albeit not statistically significant in all examined datasets, tends to have longer overall survival in both LGG and GBM samples (Fig 3C-F). Obviously, *PRKCG* expression has the potential capability to differentiate samples with diverse survival states, which would be of critical significance for accurate glioma subtyping, better therapeutic decisions and precision healthcare.

**Fig 3.**
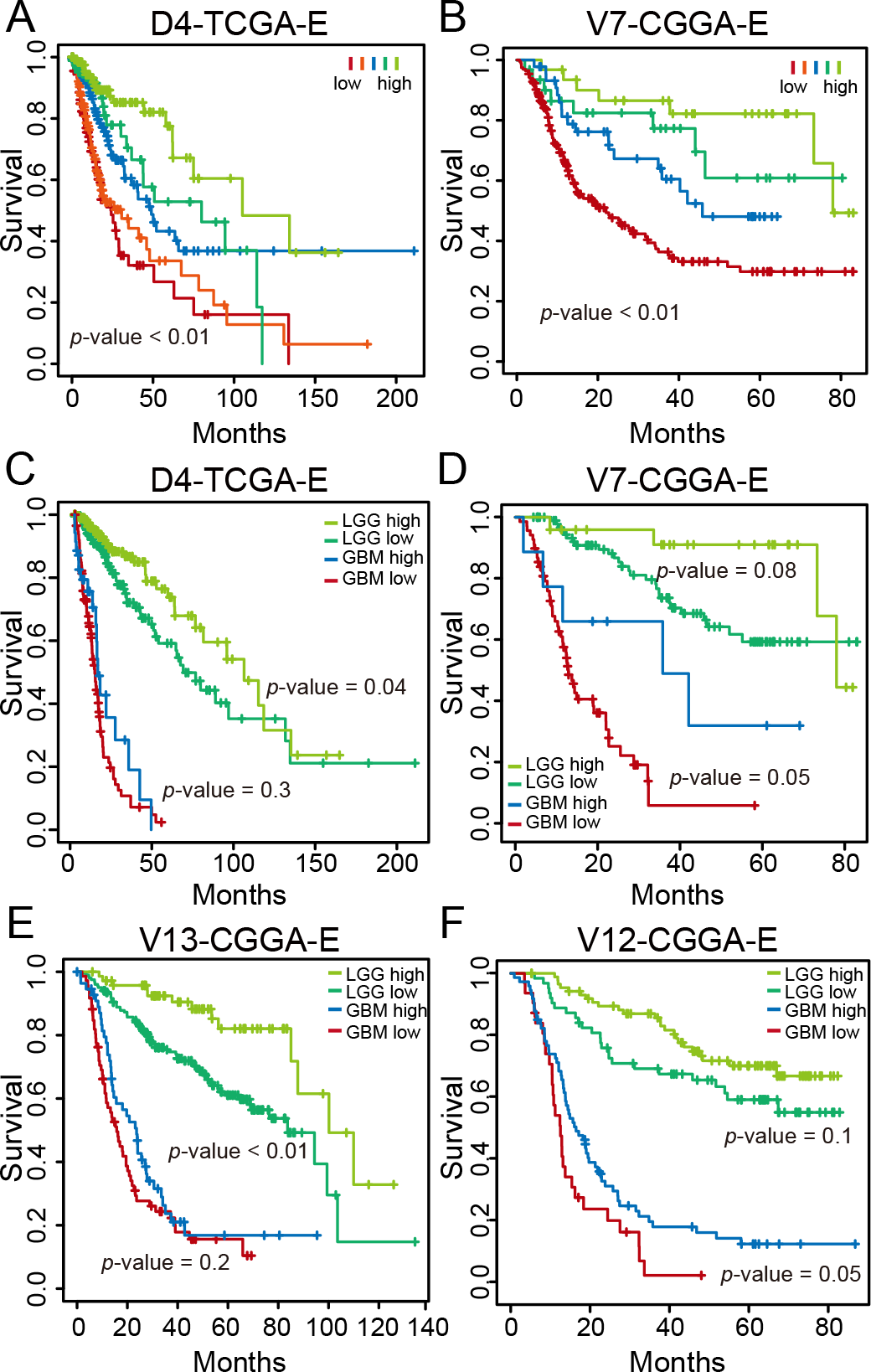
Expression profiles of *PRKCG* associated with survival. Glioma samples were divided into different groups based on *PRKCG* expression (panels A and B). LGG and GBM samples were divided into two groups with high expression and low expression, respectively (panels C to F). All these datasets can be publicly accessible at ftp://download.big.ac.cn/glioma_data/. The log-rank tests were used to examine the statistical significance between different survival curves.

### *PRKCG* is significantly differentially methylated among normal, LGG and GBM samples

Since *PRKCG* harbors two CpG sites (namely, cg26626089 and cg04518808) that are located in the promoter region and covered in both HumanMethylation27 (27K) and HumanMethylation450 (450K) BeadChip datasets, we then systematically investigated DNA methylation profiles of these two sites among normal, LGG and GBM samples. Apparently, the two sites show hypermethylation in GBM patients compared with normal samples (Fig 4), which is more significant for cg26626089 (Fig 4A and 4C; *p-*value < 0.01, Wilcoxon test). Furthermore, we examined the variation of methylation level by using whole-genome bisulfite sequencing data of six GBM samples from TCGA and one normal sample from UCSC (2017 version; http://genome.ucsc.edu, last accessed on 12 May 2019). Consistently, most GBM patients show higher methylation levels than normal samples (S5 Fig). In addition, considering different-grade gliomas, both sites present much lower methylation levels in GBM samples than LGG samples (Fig 4E-H; *p-*value < 0.01, Wilcoxon test). We further investigated its methylation in pGBM and rGBM and obtained contradictory results in different populations; *PRKCG* methylation shows no significant difference in the Chinese population (V10-CGGA-M) (S6A and S6B Fig; *p-*value > 0.05, Wilcoxon test) but significantly difference in the Switzerland population (V14-GSE60274-M) (70) (S6C and S6D Fig; *p-*value < 0.05, Wilcoxon test). This is most likely caused by the population genetic difference and/or the small sample size (both datasets have < 5 rGBM samples).

**Fig 4.**
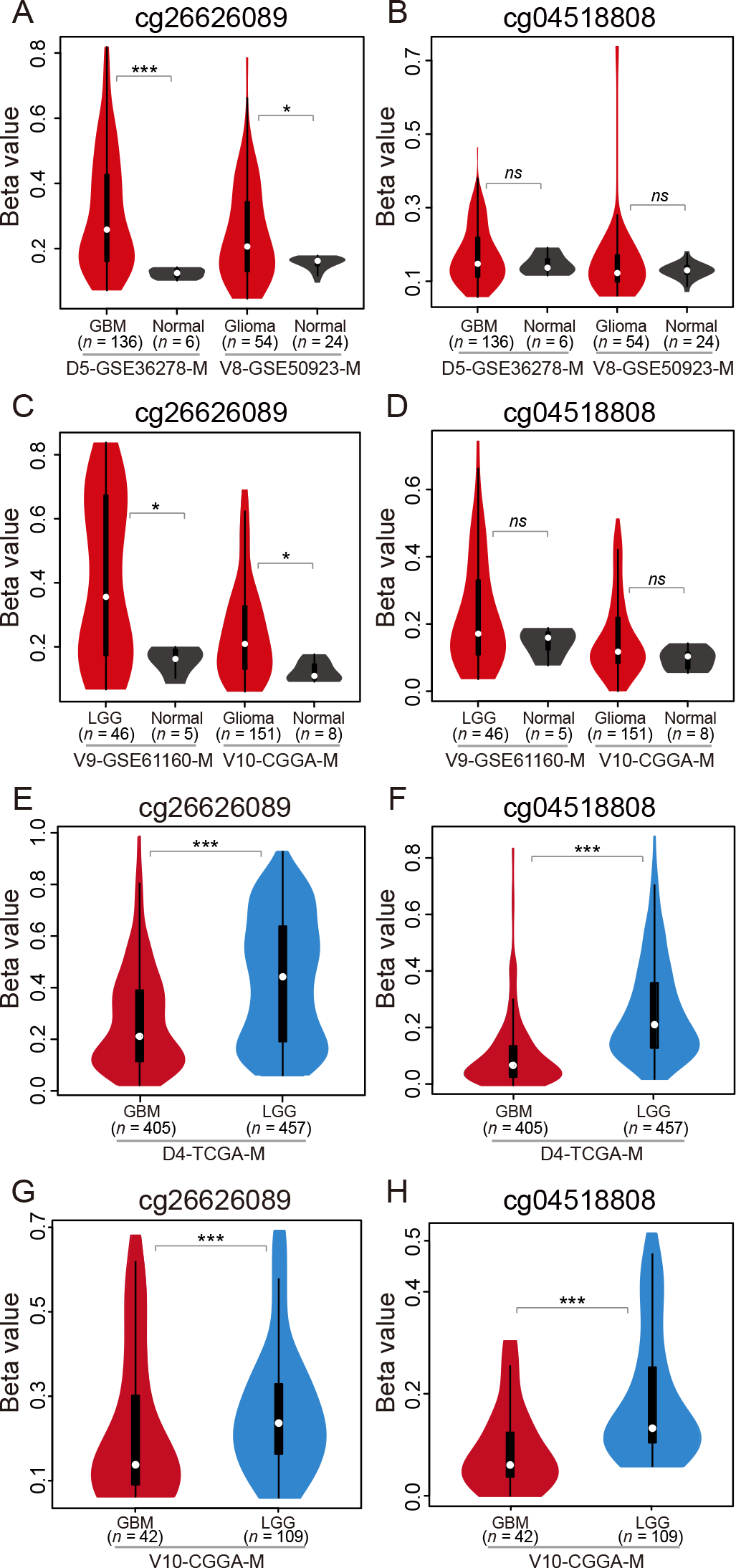
DNA methylation profiles across normal, LGG and GBM samples. *PRKCG* methylation profiles were compared between GBM and normal samples (panels A to D), and between LGG and GBM samples (panels E to H). All these datasets can be publicly accessible at ftp://download.big.ac.cn/glioma_data/. The Wilcoxon tests were used and their statistical significance levels were coded by: *ns p*>0.05, * *p*<0.05, ** *p*<0.01 and *** *p*<0.001.

Collectively, *PRKCG* is significantly differentially expressed/methylated among normal, LGG and GBM samples. Compared with normal samples, *PRKCG* presents lower expression and higher methylation in glioma samples. With tumor malignancy, *PRKCG* methylation and expression are both on the decrease (discussed below).

### Combined methylation signatures of *PRKCG* and *MGMT* are more effective in treatment prediction

It is known that *MGMT* encodes a DNA-repair protein and hypermethylation of *MGMT* is associated with diminished DNA-repair activity, accordingly allowing the alkylating drug temozolomide (TMZ) to have greater effect in GBM treatment (71, 72). In our study, we obtained consistent results that patients with methylated *MGMT* are more sensitive to TMZ treatment than those with unmethylated *MGMT* (Fig 5A; *p*-value < 0.01, log-rank test).

**Fig 5.**
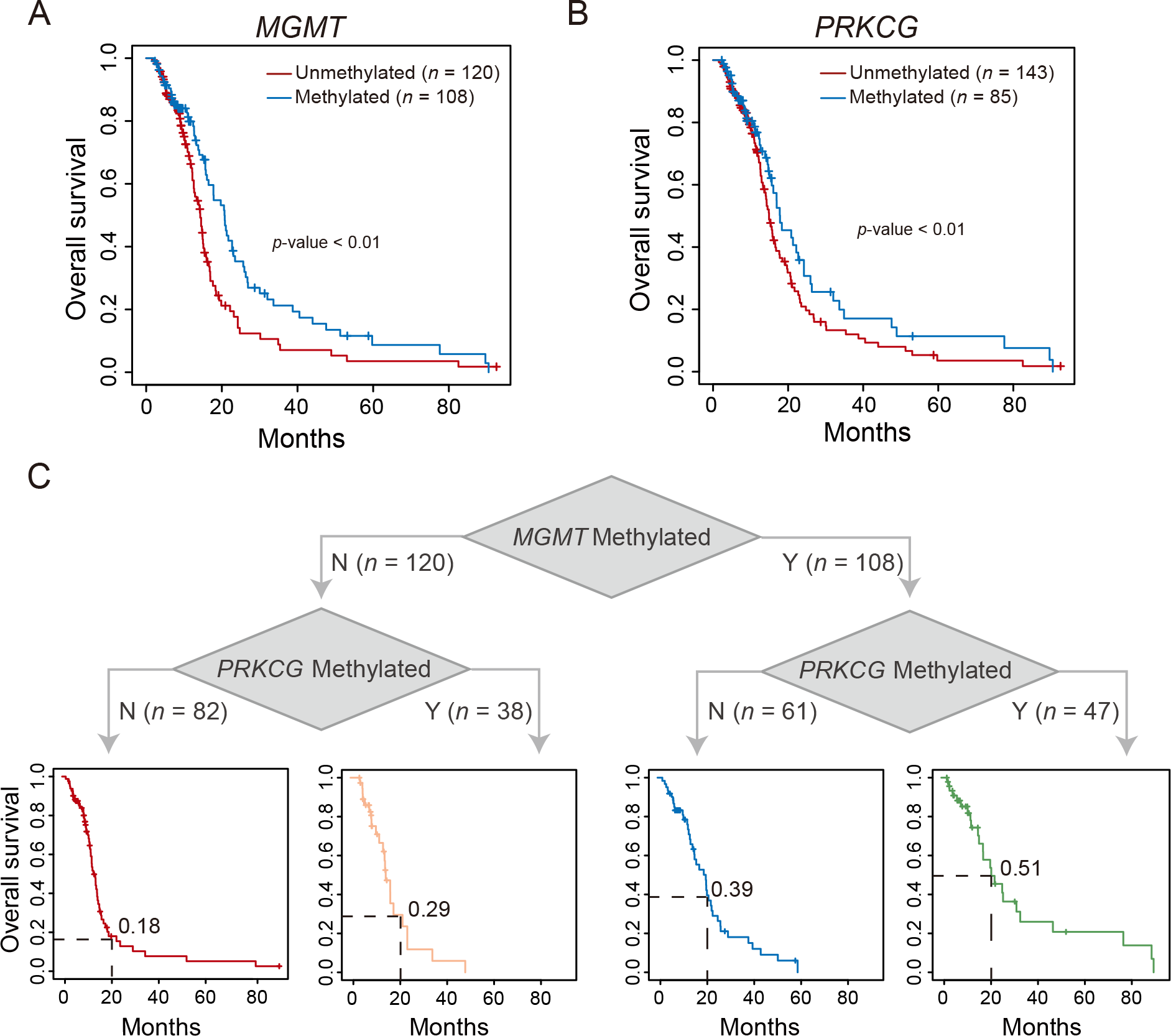
Combined DNA methylation signatures of *MGMT* and *PRKCG* for treatment prediction. (A) Kaplan-Meier survival curves for GBM patients with TMZ treatment based on *MGMT* methylation. **(B)** Kaplan-Meier survival curves for GBM patients with TMZ treatment based on *PRKCG* (cg26626089) methylation. **(C)** Kaplan-Meier survival curves for GBM patients with TMZ treatment based on *MGMT* and *PRKCG* combined methylation signatures.

Considering that a single molecular biomarker might be lack of sufficient prediction power and thus fail to determine the clinical therapeutic efficacy due to tumor heterogeneity (73), we sought to examine the predictive potential of *PRKCG* for TMZ using 228 GBM samples with matched DNA methylation and clinical data. We discovered that among the two CpG sites of *PRKCG* (cg26626089 and cg04518808), cg26626089 is able to classify patients into two groups with distinct survival advantages, as patients with methylated cg26626089 have significantly longer survival than those with unmethylated cg26626089 (Fig 5B and S7A Fig). By combining *PRKCG* (cg26626089) with *MGMT*, intriguingly, GBM patients receiving TMZ treatment can be classified into four groups that exhibit significantly different survivals (Fig 5C and S7B Fig; *p*-value < 0.01, log-rank test). The four groups, namely, *MGMT*-unmethylated + *PRKCG*-unmethylated, *MGMT*-unmethylated + *PRKCG*-methylated, *MGMT*-methylated + *PRKCG*-unmethylated, and *MGMT*-methylated + *PRKCG*-methylated, present gradually improved longer survivals, as their 20-month OS rates are 0.18, 0.29, 0.39 and 0.51 (Fig 5C), respectively, implying that the combined methylation signatures of *PRKCG* and *MGMT* might guide more accurate GBM stratification and achieve better personalized therapeutic decisions. Noticeably, elevated *MGMT* expression is associated with TMZ resistance (33). Similarly, we consistently detected that the both-methylated group with better survival shows significantly lower expression of *MGMT* (S7C Fig).

### Multi-omics molecular profiles of *PRKCG*

Based on multi-omics profiles of *PRKCG*, we explored the relationship between *PRKCG* and classical molecular features/glioma grades. First, *PRKCG* is located on the chromosome 19q13.42, unifying previous findings that 1p/19q codel is closely associated with glioma. Consistently, *PRKCG* CNV is closely associated with 19q status (Fig 6A and S8 Fig). Second, *PRKCG* methylation (cg26626089) is associated with *IDH* status, agreeing well with the finding that *IDH*-mut is an initiating event that remodels the glioma methylome, resulting in extensive DNA hypermethylation (12, 74) and thus most likely indicating that *PRKCG* methylation is a passenger of *IDH-*mut status. As LGG samples are always associated with *IDH*-mut and GBM samples are associated with *IDH*-WT, it is not difficult to understand why the methylation level of *PRKCG* is significantly lower in GBM than in LGG (Fig 4E-H). At the same time, such higher expression level and higher methylation level lead to the suspicion whether *PRKCG* expression in glioma is positively regulated by its DNA methylation or is attributable to its CNV.

**Fig 6.**
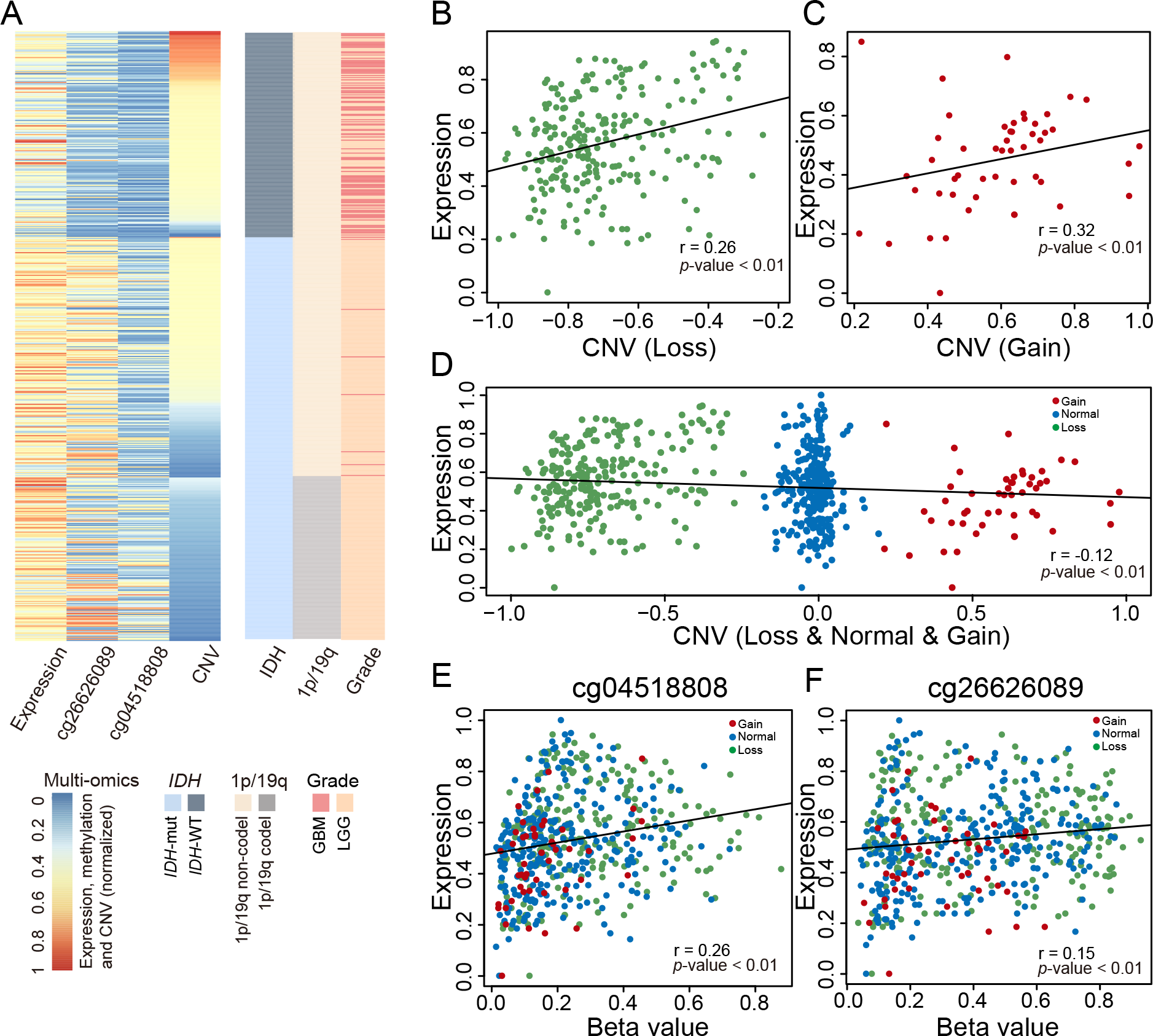
Multi-omics molecular profiles of *PRKCG*. (A) Association of *PRKCG*’s multi-omics signatures with *IDH*, 1p/19q status and WHO grade. **(B)** Correlation between *PRKCG* expression and CNV Loss. (**C**) Correlation between *PRKCG* expression and CNV Gain. (**D**) Correlation between *PRKCG* expression and all CNV status. **(E)** Correlation between *PRKCG* expression and DNA methylation of the CpG site cg04518808. **(F)** Correlation between *PRKCG* expression and DNA methylation of the CpG site cg26626089.

As expected, when considering CNV loss and gain separately, *PRKCG* CNV is positively correlated with its expression, as observed in the CNV loss and gain groups, respectively (Fig 6B and 6C; *p*-value < 0.01, Spearman correlation = 0.26/0.32). However, such positive correlation is absent when ignoring the difference of CNV status (Fig 6D). According to the dosage effect theory (75), the CNV loss group should not express more *PRKCG* than the CNV gain group. This implies that there is probably another factor rather than CNV to dominantly regulate *PRKCG* expression in glioma. Although it contradicts the commonly accepted negative association between gene expression and promoter CpG methylation, a large-scale pan-cancer analysis has also revealed a positive correlation between promoter CpG methylation and gene expression (76). Consistently, we did observe significant positive correlations between *PRKCG* expression and CpG methylation within the promoter region (Fig 6E and 6F). This positive regulation of CpG methylation is quite strong, which significantly improves *PRKCG* expression in LGG samples; even these samples exhibit obvious CNV loss (Fig 6A). Thus, it is presumably suggested that *PRKCG* is most likely regulated in different ways by DNA methylation, which negatively regulates *PRKCG* expression from normal to tumor but positively regulates the expression within tumor. However, gene expression regulation is a more complicated process involving multiple factors aside from DNA methylation and CNV and more efforts on comprehensive and in-depth molecular characterization of glioma are highly needed to elucidate glioma pathogenesis.

### *PRKCG*-like genes may present heterogeneous roles in glioma tumorigenesis

To better understand the regulation pattern of *PRKCG*, we further identified a total of 542 *PRKCG*-like genes that possess expression and DNA methylation patterns similar with *PRKCG* (see Materials and Methods) and investigated whether these genes present heterogeneous roles in glioma tumorigenesis (Fig 7A-B and S3 Table). Noticeably, some of these *PRKCG*-like genes have already been reported to be closely related with glioma (77–81). For instance, *AKAP6* (A-kinase anchoring protein 6) polymorphisms are associated with glioma susceptibility and prognosis (80); Phosphorylated *SATB1* (SATB homeobox 1) contributes to the invasive and proliferative of GBM cells and is associated with glioma prognosis (77); Higher expression of *CDK17* (cyclin dependent kinase 17) is indicative of longer overall survival (78); *PTPRM* (protein tyrosine phosphatase receptor type M) expression is significantly reduced in GBM by contrast to LGG and higher expression is indicative of longer overall survival (79); and *CHD5* (chromodomain helicase DNA binding protein 5) might act as a tumor suppressor and its lower expression is associated with poor prognosis in glioma (81). Among these *PRKCG*-like genes, we revealed that 114 genes show negative correlations between methylation level and expression level, whereas 297 genes present positive correlations (S3 Table). We further performed gene set enrichment analysis for these two groups’ genes. We found that the negatively-correlated genes are primarily enriched in the MAPK and cGMP-PKG signaling pathways, which are essential for tumor cell proliferation and differentiation (82, 83). On the contrary, the positively-correlated genes are significantly involved in pathways relevant to cancer, inflammation and nerve synapse (Fig 7C). Thus, *PRKCG*-like genes that exhibit the positive correlation between DNA methylation and expression, presumably present heterogeneous roles in glioma tumorigenesis with complex molecular mechanisms that need further extensive explorations both bioinformatically and experimentally.

**Fig 7.**
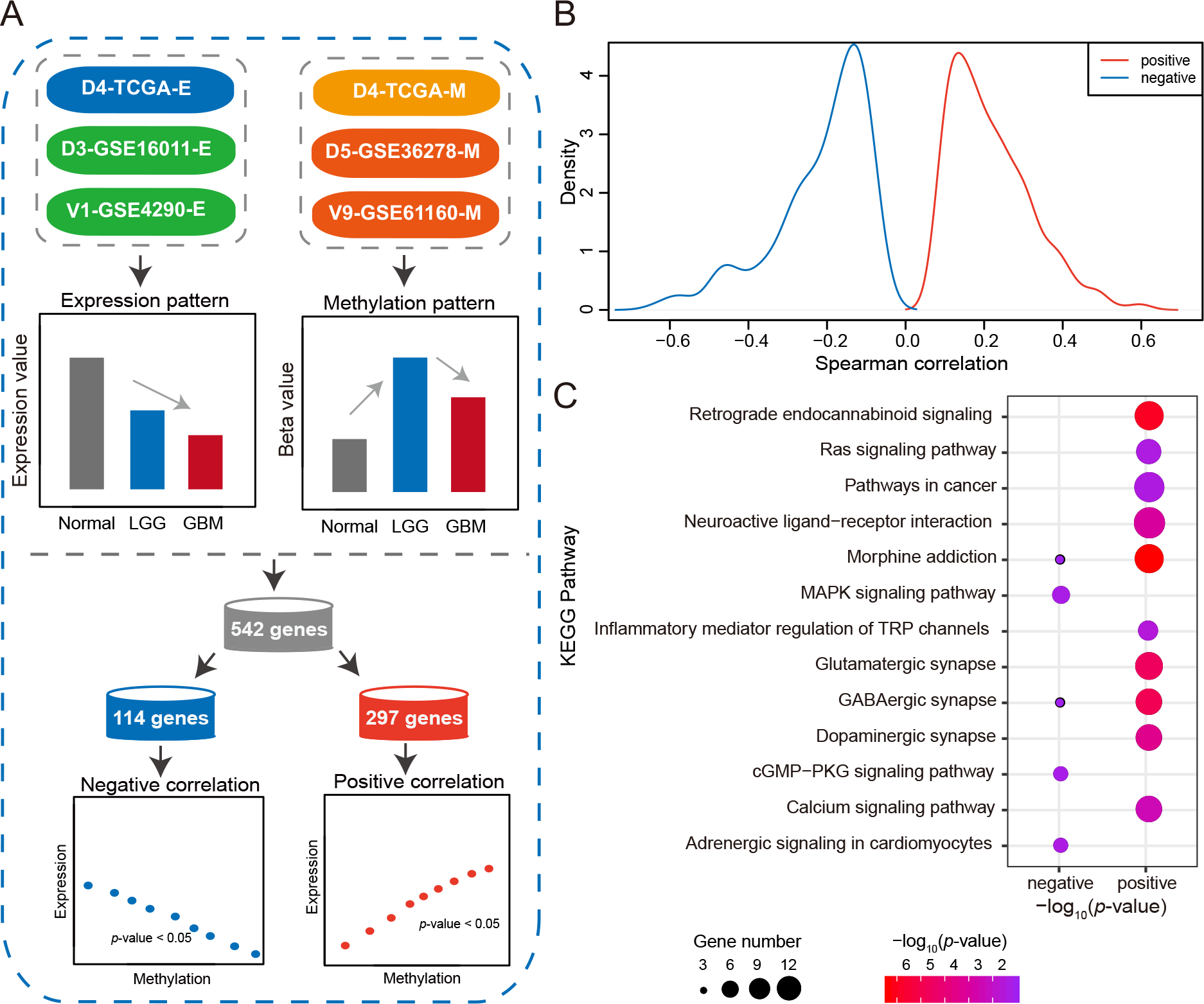
Identification and characterization of *PRKCG*-like genes. (A) Identification of *PRKCG*-like genes, including two types of genes that possess negative and positive correlations between expression and DNA methylation, respectively. **(B)** Spearman correlations of two types of *PRKCG*-like genes. **(C)** The KEGG pathway enrichment of two types of *PRKCG*-like genes.

## Conclusion

The rapid advancement of sequencing technologies enables large amounts of glioma data generated from different projects and studies throughout the world. Therefore, it has become crucially significant on how to make full use of these valuable data for computational identification and characterization of glioma candidate biomarkers. In this study, we, for the first time, assembled the most comprehensive collection of public glioma datasets with multi-omics data types and different populations/countries. Through comprehensive molecular profiling, we identified that *PRKCG*, a brain-specific gene detectable in CSF, is a potential biomarker for glioma diagnosis, prognosis and treatment prediction. Specifically, it presents lower expression and higher methylation in glioma samples than normal samples. *PRKCG* expression/methylation change from high to low is indicative of glioma progression from low-grade to high-grade and high RNA expression is suggestive of good survival. Importantly, *PRKCG* in combination with *MGMT* is more effective to yield precise survival outcomes after TMZ chemotherapy. In harmony with classical biomarkers, *PRKCG* as well as *PRKCG*-like genes may play important and heterogeneous roles in glioma tumorigenesis. In the era of big data, our findings highlight the importance of computational integrative multi-omics profiling and represent a data-driven scheme toward precision tumor subtyping, accurate therapeutic decisions and better personalized healthcare.

## Acknowledgements

We thank Jun Yu, Songnian Hu, Fangqing Zhao, Yu Xue, Zheng Zhao, Jiabao Cao, Jian Sang, Guangyi Niu, Man Li, and Yang Zhang for their valuable comments and discussions on this work.

## Author Contributions

Conceptualization, L.L. and W.G.Y.; Methodology, L.L., W.G.Y., and Z.Z.; Validation, L.L. and W.L.G.; Formal Analysis, L.L., W.G.Y., W.L.G., Y.C.L. and L.M.W.; Investigation, L.L. and W.G.Y.; Writing – Original Draft, L.L.; Writing – Review & Editing, M.L.N., W.L.G., S.S.H., H.L.L., and Z.Z.; Funding Acquisition, M.L.N. and Z.Z.; Supervision, M.L.N. and Z.Z.

## Supporting information captions

**S1 Fig.**
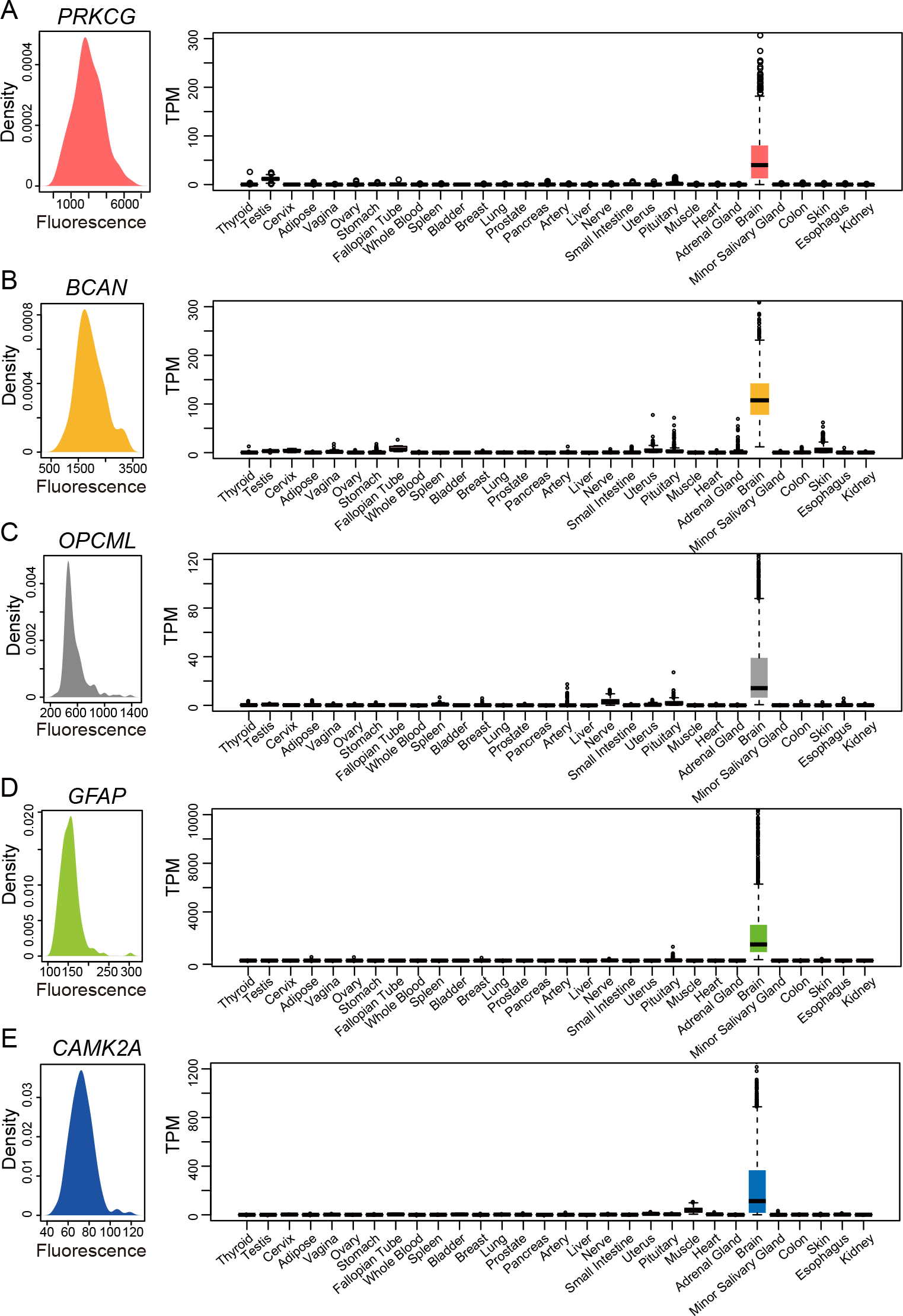
Protein level distributions in CSF and RNA expression profiles of *PRKCG* (A), ***BCAN* (B), *OPCML* (C), *GFAP* (D), *CAMK2A* (E) across 30 normal human tissues.**

**S2 Fig.**
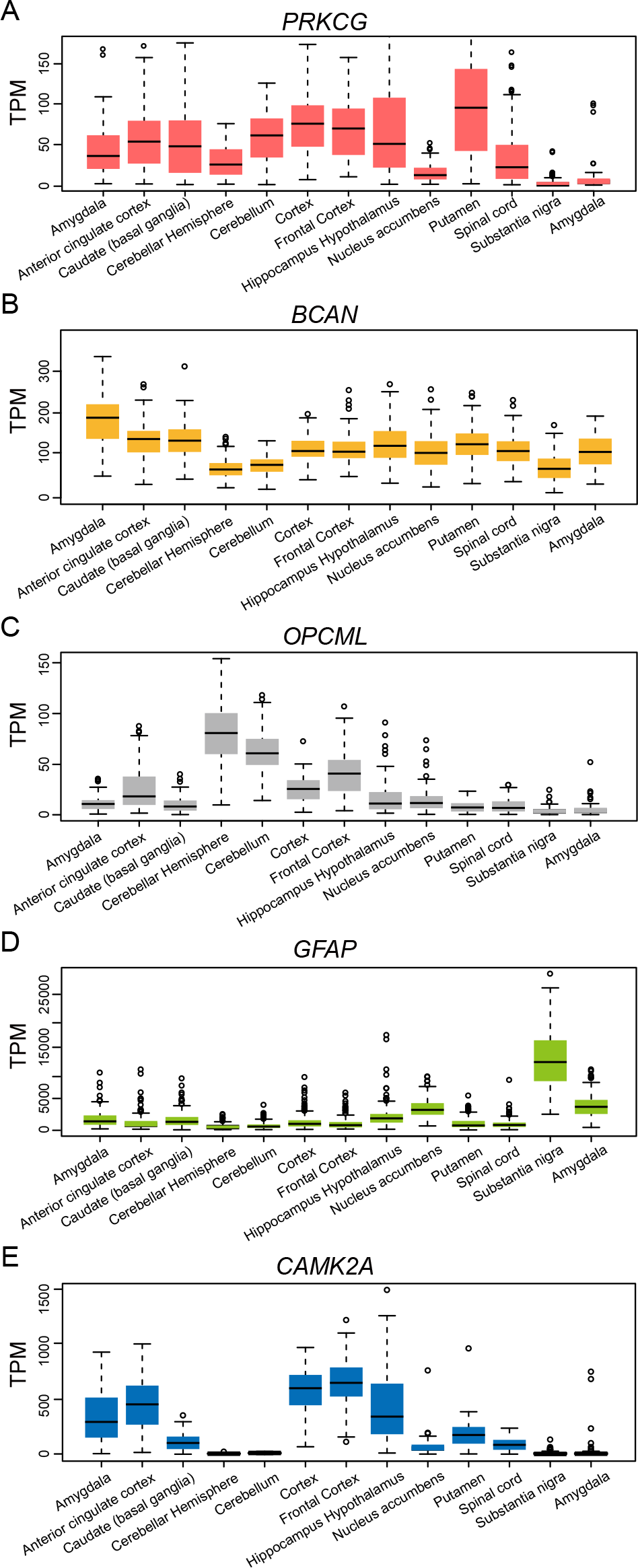
Expression profiles of *PRKCG* (A), *BCAN* (B), *OPCML* (C), *GFAP* (D), and *CAMK2A* (E) across 13 human brain regions.

**S3 Fig.**
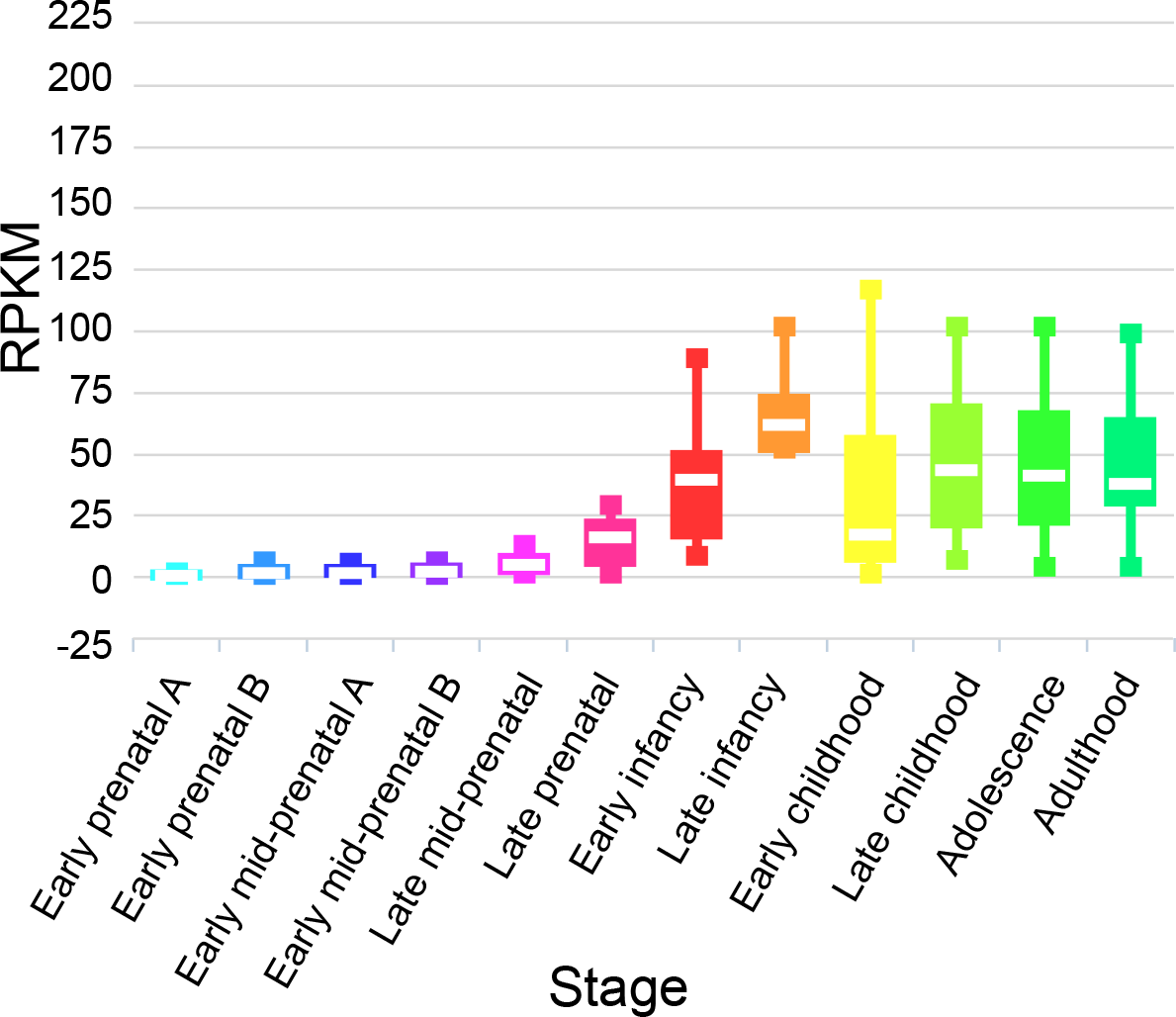
Expression profiles of *PRKCG* in brain developmental stages.

**S4 Fig.**
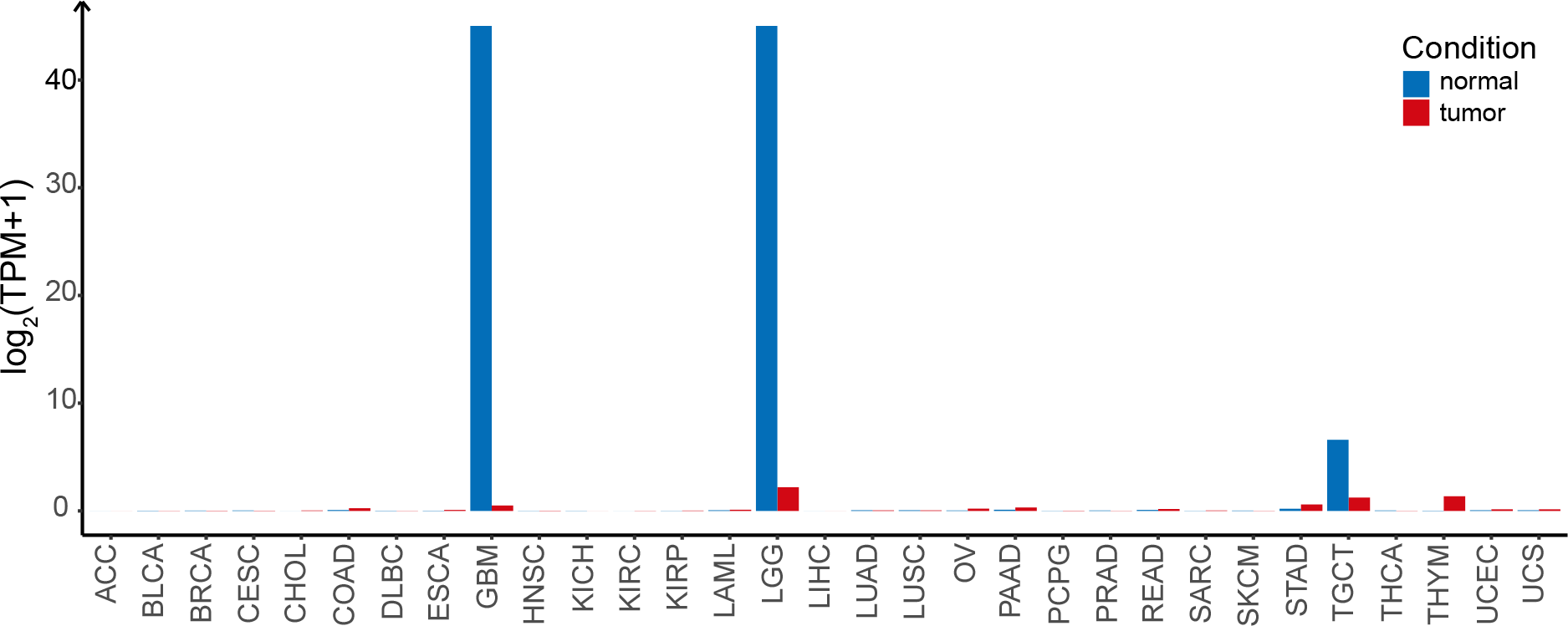
Expression profiles of *PRKCG* across 31 human tumor and normal tissues.

**S5 Fig.**
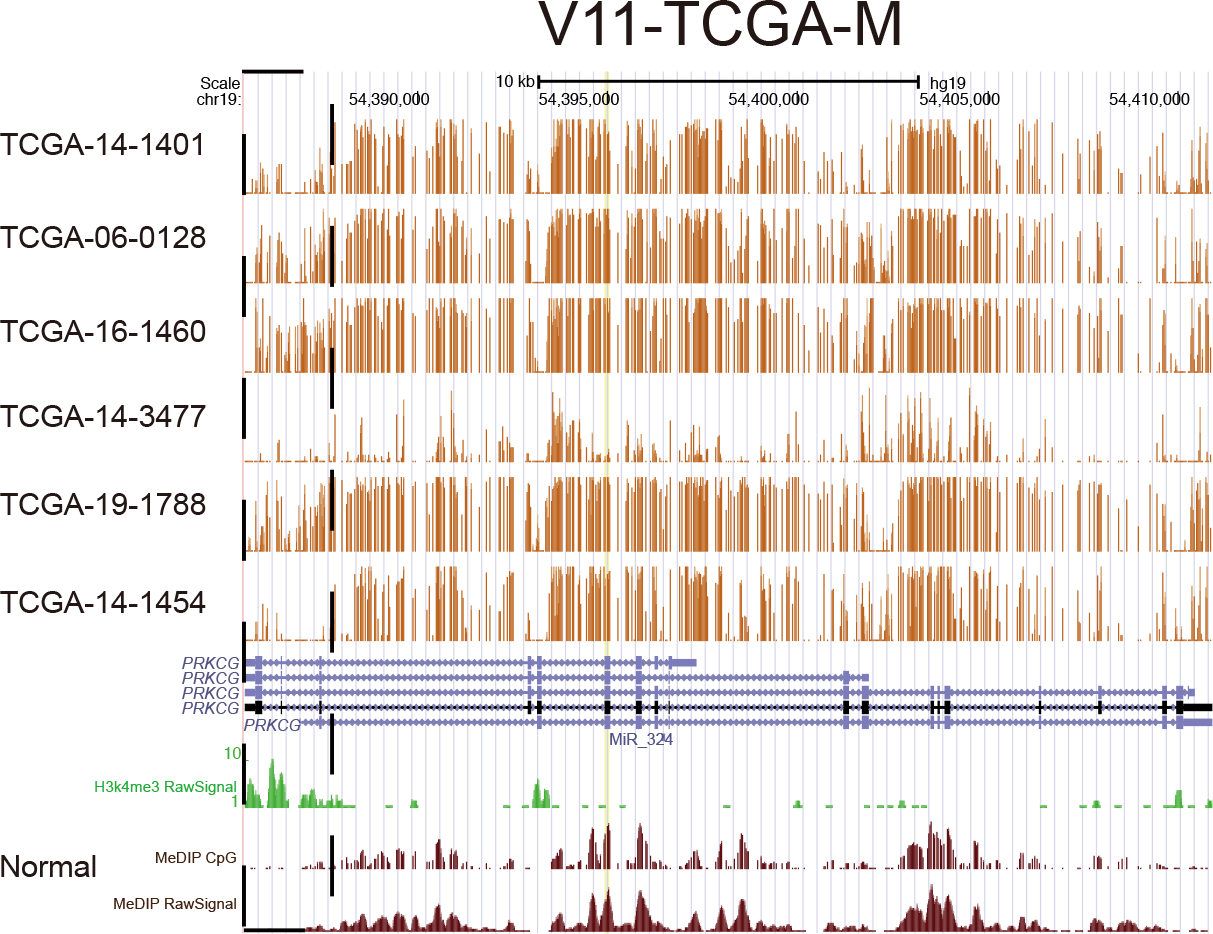
Bisulfite DNA methylation profiles of *PRKCG* across six GBM samples and one normal sample.

**S6 Fig.**
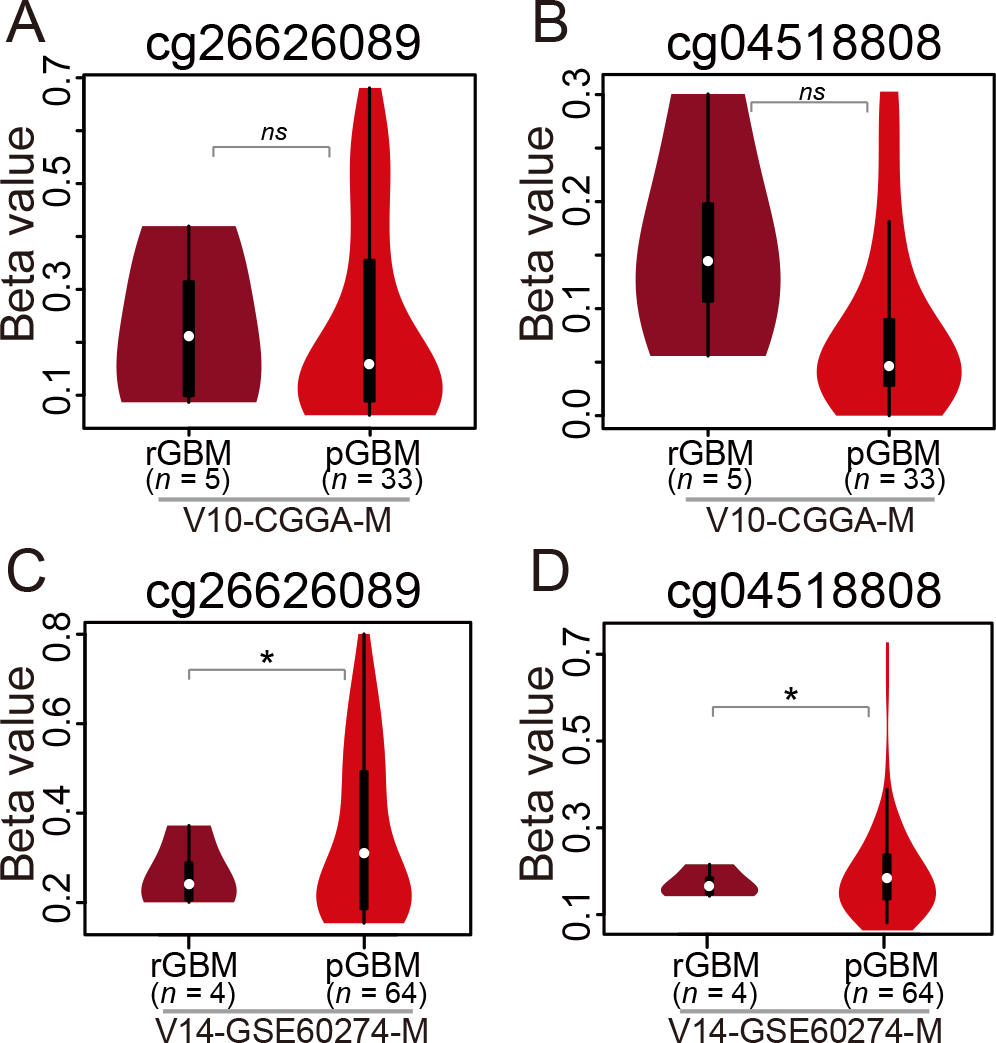
DNA methylation profiles of *PRKCG* in recurrent GBM (rGBM) and primary GBM (pGBM) samples. *PRKCG* methylation profiles were compared between rGBM and pGBM samples (V10-CGGA-M in panels A and B and V14-GSE60274-M in panels C and D). All these datasets can be publicly accessible at ftp://download.big.ac.cn/glioma_data/. The Wilcoxon tests were used and the statistical significance levels were coded by: *ns p*>0.05, * *p*<0.05, ** *p*<0.01 and *** *p*<0.001.

**S7 Fig.**
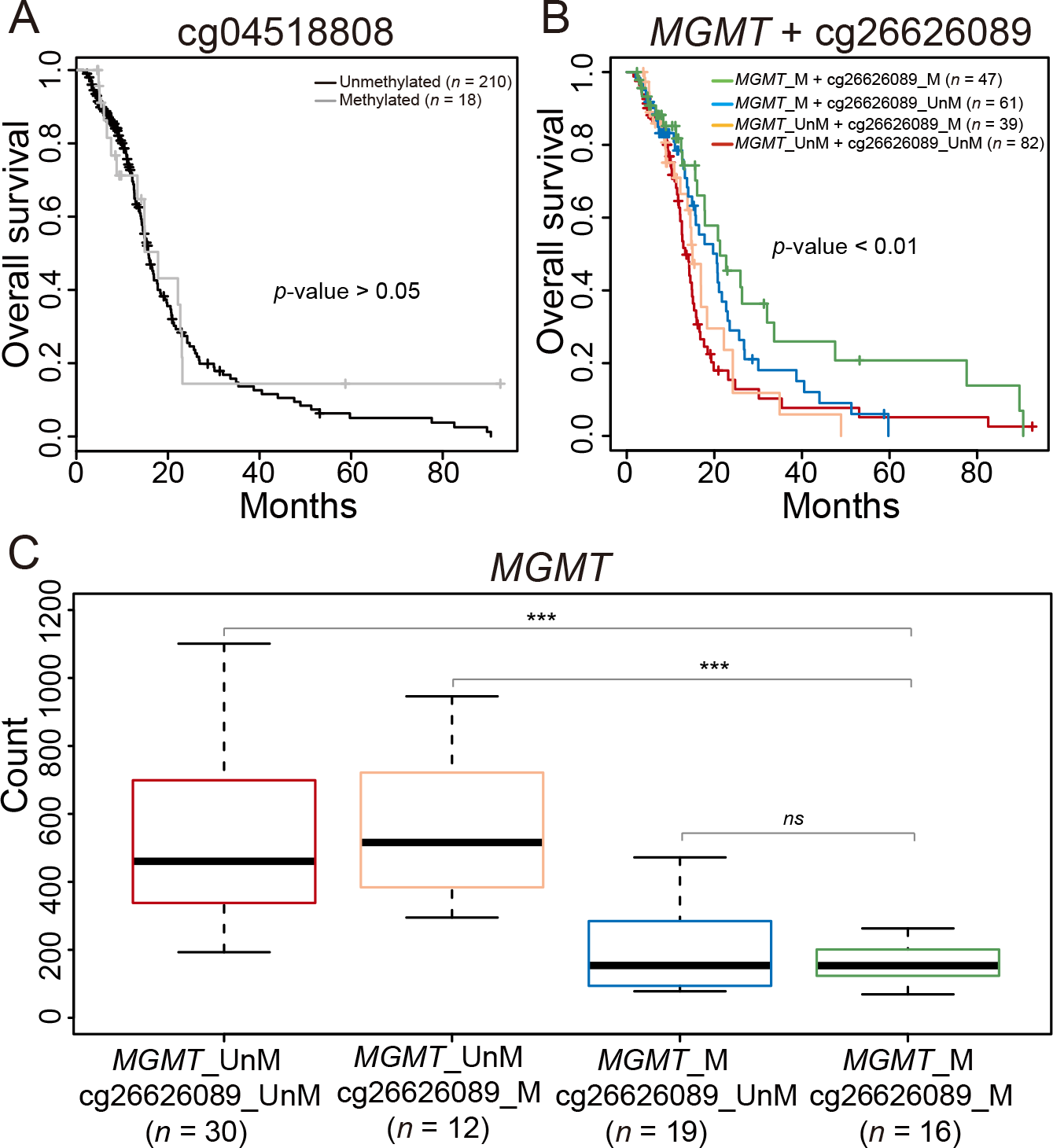
**Predictive potential of *PRKCG* DNA methylation.** (**A**) Kaplan-Meier survival curves for GBM patients with TMZ treatment based on *PRKCG* (cg04518808) methylation. (**B**) Methylation site cg26626089 in combination with *MGMT*, which were used to classify GBM patients into four groups. (**C**) Expression profiles of *MGMT* in the four groups.

**S8 Fig.**
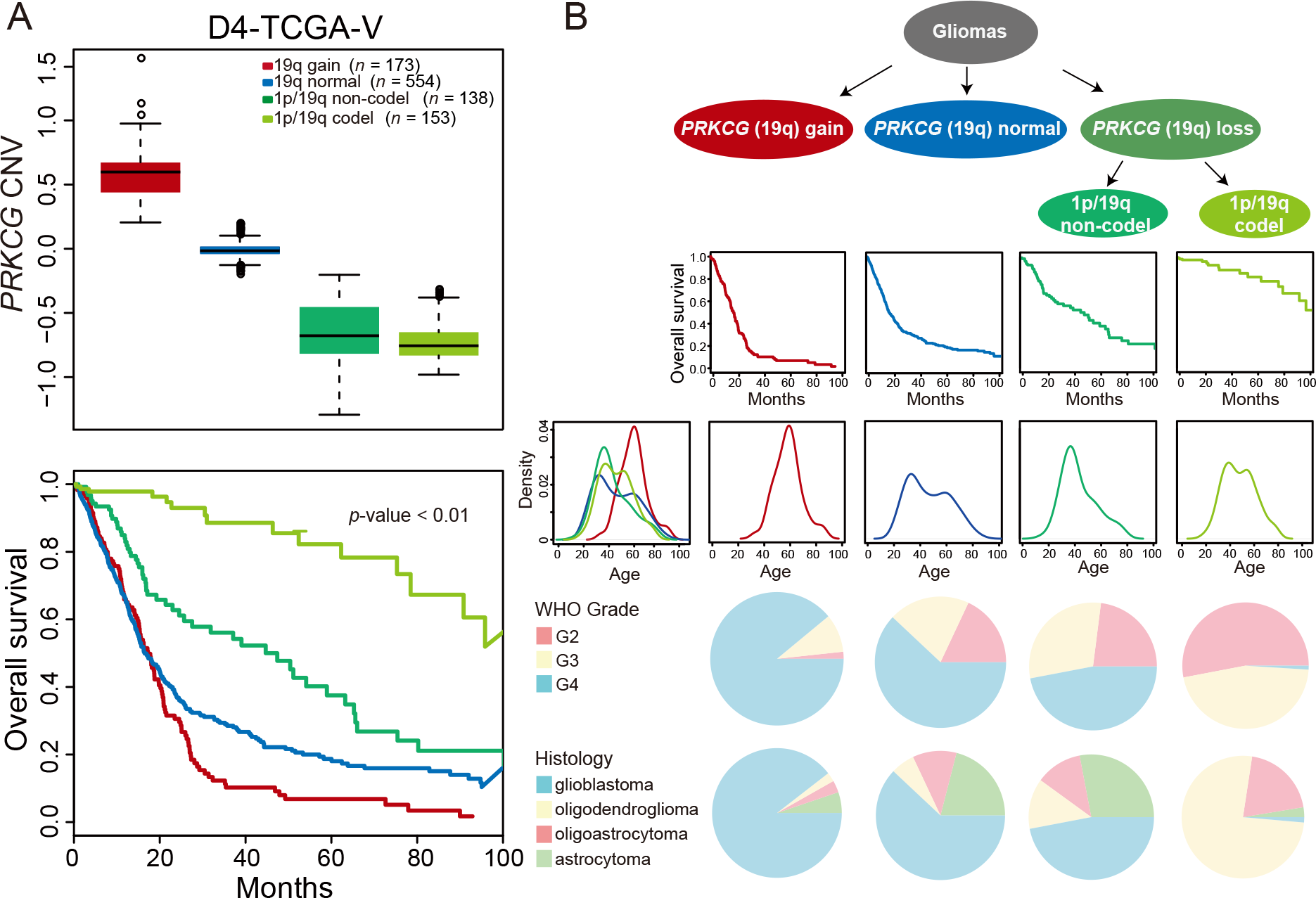
*PRKCG* CNV associated with survival. (A) Four groups of glioma samples were divided based on the 1p/19q status (19q gain, 19q normal, 1p/19q non-codel, and 1p/19q codel). **(B)** Kaplan-Meier survival probability, age, WHO grade and histology of the four groups.

**S1 Table.** 15,176 genes’ RNA expression levels across 30 normal human tissues.

**S2 Table.** Tissue specificity τ values and maximum expression levels of the top 100 brain-specific protein-coding genes.

**S3 Table.** *PRKCG*-like genes and their annotations.

## References

1. Schwartzbaum JA, Fisher JL, Aldape KD, Wrensch M. Epidemiology and molecular pathology of glioma. Nat Clin Pract Neurol. 2006;2(9):494–503; quiz 1 p following 16.

2. Maher EA, Furnari FB, Bachoo RM, Rowitch DH, Louis DN, Cavenee WK, et al. Malignant glioma: genetics and biology of a grave matter. Genes Dev. 2001;15(11):1311–33.

3. Stewart LA. Chemotherapy in adult high-grade glioma: a systematic review and meta-analysis of individual patient data from 12 randomised trials. Lancet. 2002;359(9311):1011–8.

4. Louis DN, Ohgaki H, Wiestler OD, Cavenee WK, Burger PC, Jouvet A, et al. The 2007 WHO classification of tumours of the central nervous system. Acta Neuropathol. 2007;114(2):97–109.

5. Wesseling P, Kros JM, Jeuken JWM. The pathological diagnosis of diffuse gliomas: towards a smart synthesis of microscopic and molecular information in a multidisciplinary context. Diagnostic Histopathology. 2011;17(11):486–94.

6. Kim YW, Koul D, Kim SH, Lucio-Eterovic AK, Freire PR, Yao J, et al. Identification of prognostic gene signatures of glioblastoma: a study based on TCGA data analysis. Neuro Oncol. 2013;15(7):829–39.

7. Wiestler B, Capper D, Holland-Letz T, Korshunov A, von Deimling A, Pfister SM, et al. ATRX loss refines the classification of anaplastic gliomas and identifies a subgroup of IDH mutant astrocytic tumors with better prognosis. Acta Neuropathol. 2013;126(3):443–51.

8. Cohen AL, Holmen SL, Colman H. IDH1 and IDH2 mutations in gliomas. Curr Neurol Neurosci Rep. 2013;13(5):345.

9. Eckel-Passow JE, Lachance DH, Molinaro AM, Walsh KM, Decker PA, Sicotte H, et al. Glioma Groups Based on 1p/19q, IDH, and TERT Promoter Mutations in Tumors. N Engl J Med. 2015;372(26):2499–508.

10. Waitkus MS, Diplas BH, Yan H. Biological Role and Therapeutic Potential of IDH Mutations in Cancer. Cancer Cell. 2018;34(2):186–95.

11. Kloosterhof NK, Bralten LBC, Dubbink HJ, French PJ, van den Bent MJ. Isocitrate dehydrogenase-1 mutations: a fundamentally new understanding of diffuse glioma? The Lancet Oncology. 2011;12(1):83–91.

12. Turcan S, Rohle D, Goenka A, Walsh LA, Fang F, Yilmaz E, et al. IDH1 mutation is sufficient to establish the glioma hypermethylator phenotype. Nature. 2012;483(7390):479–83.

13. Jenkins RB, Blair H, Ballman KV, Giannini C, Arusell RM, Law M, et al. A t(1;19)(q10;p10) mediates the combined deletions of 1p and 19q and predicts a better prognosis of patients with oligodendroglioma. Cancer Res. 2006;66(20):9852–61.

14. Cairncross G, Wang M, Shaw E, Jenkins R, Brachman D, Buckner J, et al. Phase III trial of chemoradiotherapy for anaplastic oligodendroglioma: long-term results of RTOG 9402. J Clin Oncol. 2013;31(3):337–43.

15. Mur P, Mollejo M, Hernandez-Iglesias T, de Lope AR, Castresana JS, Garcia JF, et al. Molecular classification defines 4 prognostically distinct glioma groups irrespective of diagnosis and grade. J Neuropathol Exp Neurol. 2015;74(3):241–9.

16. Molinaro AM, Taylor JW, Wiencke JK, Wrensch MR. Genetic and molecular epidemiology of adult diffuse glioma. Nat Rev Neurol. 2019;15(7):405–17.

17. Louis DN, Perry A, Reifenberger G, von Deimling A, Figarella-Branger D, Cavenee WK, et al. The 2016 World Health Organization Classification of Tumors of the Central Nervous System: a summary. Acta Neuropathol. 2016;131(6):803–20.

18. Flynn JR, Wang L, Gillespie DL, Stoddard GJ, Reid JK, Owens J, et al. Hypoxia-regulated protein expression, patient characteristics, and preoperative imaging as predictors of survival in adults with glioblastoma multiforme. Cancer. 2008;113(5):1032–42.

19. Verhaak RGW, Hoadley KA, Purdom E, Wang V, Qi Y, Wilkerson MD, et al. Integrated Genomic Analysis Identifies Clinically Relevant Subtypes of Glioblastoma Characterized by Abnormalities in PDGFRA, IDH1, EGFR, and NF1. Cancer Cell. 2010;17(1):98–110.

20. Wang Q, Hu B, Hu X, Kim H, Squatrito M, Scarpace L, et al. Tumor evolution of glioma intrinsic gene expression subtype associates with immunological changes in the microenvironment. Cancer Cell. 2017;32(1):42–56.

21. Gao W-Z, Guo L-M, Xu T-Q, Yin Y-H, Jia F. Identification of a multidimensional transcriptome signature for survival prediction of postoperative glioblastoma multiforme patients. Journal of Translational Medicine. 2018;16(1).

22. Hatanpaa KJ, Burma S, Zhao D, Habib AA. Epidermal growth factor receptor in glioma: signal transduction, neuropathology, imaging, and radioresistance. Neoplasia. 2010;12(9):675–84.

23. Gan HK, Kaye AH, Luwor RB. The EGFRvIII variant in glioblastoma multiforme. J Clin Neurosci. 2009;16(6):748–54.

24. Mukherjee B, McEllin B, Camacho CV, Tomimatsu N, Sirasanagandala S, Nannepaga S, et al. EGFRvIII and DNA Double-Strand Break Repair: A Molecular Mechanism for Radioresistance in Glioblastoma. Cancer Research. 2009;69(10):4252–9.

25. Zhang S, Zhao BS, Zhou A, Lin K, Zheng S, Lu Z, et al. m(6)A Demethylase ALKBH5 Maintains Tumorigenicity of Glioblastoma Stem-like Cells by Sustaining FOXM1 Expression and Cell Proliferation Program. Cancer Cell. 2017;31(4):591–606 e6.

26. Esteller M. Cancer epigenomics: DNA methylomes and histone-modification maps. Nat Rev Genet. 2007;8(4):286–98.

27. Park JY, Lee JE, Park JB, Yoo H, Lee SH, Kim JH. Roles of Long Non-Coding RNAs on Tumorigenesis and Glioma Development. Brain Tumor Res Treat. 2014;2(1):1–6.

28. Zang L, Kondengaden SM, Che F, Wang L, Heng X. Potential Epigenetic-Based Therapeutic Targets for Glioma. Front Mol Neurosci. 2018;11:408.

29. Malzkorn B, Wolter M, Riemenschneider MJ, Reifenberger G. Unraveling the Glioma Epigenome-From Molecular Mechanisms to Novel Biomarkers and Therapeutic Targets. Brain Pathology. 2011;21(6):619–32.

30. Binabaj MM, Bahrami A, ShahidSales S, Joodi M, Joudi Mashhad M, Hassanian SM, et al. The prognostic value of MGMT promoter methylation in glioblastoma: A meta-analysis of clinical trials. J Cell Physiol. 2018;233(1):378–86.

31. Wick W, Weller M, van den Bent M, Sanson M, Weiler M, von Deimling A, et al. MGMT testing--the challenges for biomarker-based glioma treatment. Nat Rev Neurol. 2014;10(7):372–85.

32. Rivera AL, Pelloski CE, Gilbert MR, Colman H, De La Cruz C, Sulman EP, et al. MGMT promoter methylation is predictive of response to radiotherapy and prognostic in the absence of adjuvant alkylating chemotherapy for glioblastoma. Neuro Oncol. 2010;12(2):116–21.

33. Hegi ME, Diserens AC, Gorlia T, Hamou MF, de Tribolet N, Weller M, et al. MGMT gene silencing and benefit from temozolomide in glioblastoma. N Engl J Med. 2005;352(10):997–1003.

34. Cancer Genome Atlas Research N, Brat DJ, Verhaak RG, Aldape KD, Yung WK, Salama SR, et al. Comprehensive, Integrative Genomic Analysis of Diffuse Lower-Grade Gliomas. N Engl J Med. 2015;372(26):2481–98.

35. Ceccarelli M, Barthel FP, Malta TM, Sabedot TS, Salama SR, Murray BA, et al. Molecular Profiling Reveals Biologically Discrete Subsets and Pathways of Progression in Diffuse Glioma. Cell. 2016;164(3):550–63.

36. Zhao Z, Zhang K, Wang Z, Wang K, Liu X, Wu F, et al. A comprehensive review of available omics data resources and molecular profiling for precision glioma studies. Biomed Rep. 2019;10(1):3–9.

37. Cao H, Wang F, Li XJ. Future Strategies on Glioma Research: From Big Data to the Clinic. Genomics Proteomics Bioinformatics. 2017;15(4):263–5.

38. Yang Y, Sui Y, Xie B, Qu H, Fang X. GliomaDB: A Web Server for Integrating Glioma Omics Data and Interactive Analysis. Genomics Proteomics Bioinformatics. 2019;17(4):465–71.

39. John Lonsdale JT, Mike Salvatore, Rebecca Phillips. The Genotype-Tissue Expression (GTEx) project. Nat Genet. 2013;45(6):580–5.

40. Puchalski. RB, Shah. N, Miller. J, Dalley. R, Nomura. SR, Yoon. J-G, et al. An anatomic transcriptional atlas of human glioblastoma. Science. 2018;360(6389):660–3.

41. Bao ZS, Chen HM, Yang MY, Zhang CB, Yu K, Ye WL, et al. RNA-seq of 272 gliomas revealed a novel, recurrent PTPRZ1-MET fusion transcript in secondary glioblastomas. Genome Res. 2014;24(11):1765–73.

42. Sun Y, Zhang W, Chen D, Lv Y, Zheng J, Lilljebjorn H, et al. A glioma classification scheme based on coexpression modules of EGFR and PDGFRA. Proc Natl Acad Sci U S A. 2014;111(9):3538–43.

43. Yanai I, Benjamin H, Shmoish M, Chalifa-Caspi V, Shklar M, Ophir R, et al. Genome-wide midrange transcription profiles reveal expression level relationships in human tissue specification. Bioinformatics. 2005;21(5):650–9.

44. Klee EW. Data mining for biomarker development: a review of tissue specificity analysis. Clin Lab Med. 2008;28(1):127–43, viii.

45. Vasmatzis G, Klee EW, Kube DM, Therneau TM, Kosari F. Quantitating tissue specificity of human genes to facilitate biomarker discovery. Bioinformatics. 2007;23(11):1348–55.

46. Kim P, Park A, Han G, Sun H, Jia P, Zhao Z. TissGDB: tissue-specific gene database in cancer. Nucleic Acids Research. 2018;46(D1):D1031–D8.

47. Prassas l, Chrystoja CC, Makawita S, Diamandis EP. Bioinformatic identification of proteins with tissue-specific expression for biomarker discovery. BMC Medicine. 2012;10(39).

48. Mohammed A, Biegert G, Adamec J, Helikar T. Identification of potential tissue-specific cancer biomarkers and development of cancer versus normal genomic classifiers. Oncotarget. 2017;8(49):85692–715.

49. Sasayama D, Hattori K, Ogawa S, Yokota Y, Matsumura R, Teraishi T, et al. Genome-wide quantitative trait loci mapping of the human cerebrospinal fluid proteome. Hum Mol Genet. 2017;26(1):44–51.

50. Gate D, Saligrama N, Leventhal O, Yang AC, Unger MS, Middeldorp J, et al. Clonally expanded CD8 T cells patrol the cerebrospinal fluid in Alzheimer’s disease. Nature. 2020.

51. Khasawneh AH, Garling RJ, Harris CA. Cerebrospinal fluid circulation: What do we know and how do we know it? Brain Circulation. 2018.

52. Sydney C. Gary CAZ, Veronica L. Chiang, Janette U. Gaw, Grace Gray, Susan Hockfield. cDNA cloning, chromosomal localization, and expression analysis of human BEHAB/brevican, a brain specific proteoglycan regulated during cortical development and in glioma. GENE. 2000;256(1-2):139–47.

53. Phillips HS, Kharbanda S, Chen R, Forrest WF, Soriano RH, Wu TD, et al. Molecular subclasses of high-grade glioma predict prognosis, delineate a pattern of disease progression, and resemble stages in neurogenesis. Cancer Cell. 2006;9(3):157–73.

54. Cook PJ, Thomas R, Kannan R, de Leon ES, Drilon A, Rosenblum MK, et al. Author Correction: Somatic chromosomal engineering identifies BCAN-NTRK1 as a potent glioma driver and therapeutic target. Nat Commun. 2018;9:16187.

55. Reed JE, Dunn JR, du Plessis DG, Shaw EJ, Reeves P, Gee AL, et al. Expression of cellular adhesion molecule ‘OPCML’ is down-regulated in gliomas and other brain tumours. Neuropathol Appl Neurobiol. 2007;33(1):77–85.

56. Carminati. PO, Mello. SS, Fachin. AL, Junta. CM, Sandrin-Garcia. P, Carlotti. CG, et al. Alterations in gene expression profiles correlated with cisplatin cytotoxicity in the glioma U343 cell line. Genetics and molecular biology. 2010;33(1):159–68.

57. Horst M, Brouwer E, Verwijnen S, Rodijk M, de Jong M, Hoeben R, et al. Targeting malignant gliomas with a glial fibrillary acidic protein (GFAP)-selective oncolytic adenovirus. J Gene Med. 2007;9(12):1071–9.

58. van Bodegraven EJ, van Asperen JV, Robe PAJ, Hol EM. Importance of GFAP isoform-specific analyses in astrocytoma. Glia. 2019;67(8):1417–33.

59. John S, Sivakumar KC, Mishra R. Bacoside A Induces Tumor Cell Death in Human Glioblastoma Cell Lines through Catastrophic Macropinocytosis. Front Mol Neurosci. 2017;10:171.

60. Long H, Liang C, Zhang X, Fang L, Wang G, Qi S, et al. Prediction and Analysis of Key Genes in Glioblastoma Based on Bioinformatics. Biomed Res Int. 2017;2017:7653101.

61. Shao Y, Chen C, Shen H, He BZ, Yu D, Jiang S, et al. GenTree, an integrated resource for analyzing the evolution and function of primate-specific coding genes. Genome Res. 2019;29(4):682–96.

62. Saito N, Shirai Y. Protein kinase C gamma (PKC gamma): function of neuron specific isotype. J Biochem. 2002;132(5):683–7.

63. Klebe S, Durr A, Rentschler A, Hahn-Barma V, Abele M, Bouslam N, et al. New mutations in protein kinase Cgamma associated with spinocerebellar ataxia type 14. Ann Neurol. 2005;58(5):720–9.

64. Yabe I, Sasaki H, Chen DH, Raskind WH, Bird TD, Yamashita I, et al. Spinocerebellar ataxia type 14 caused by a mutation in protein kinase C gamma. Arch Neurol. 2003;60(12):1749–51.

65. do Carmo A, Balca-Silva J, Matias D, Lopes MC. PKC signaling in glioblastoma. Cancer Biol Ther. 2013;14(4):287–94.

66. Reina-Campos M, Diaz-Meco MT, Moscat J. The Dual Roles of the Atypical Protein Kinase Cs in Cancer. Cancer Cell. 2019;36(3):218–35.

67. Hu L, Li X, Liu Q, Xu J, Ge H, Wang Z, et al. UBE2S, a novel substrate of Akt1, associates with Ku70 and regulates DNA repair and glioblastoma multiforme resistance to chemotherapy. Oncogene. 2017;36(8):1145–56.

68. Jung E, Osswald M, Blaes J, Wiestler B, Sahm F, Schmenger T, et al. Tweety-Homolog 1 Drives Brain Colonization of Gliomas. J Neurosci. 2017;37(29):6837–50.

69. Catriona M. Dowling SLH, James J. Phelan Mary Clare Cathcart, Stephen P. Finn Brian Mehigan Paul McCormick, John C. Coffey, Jacintha, Kiely OSaPA. Expression of protein kinase C gamma promotes cell migration in colon cancer. Oncotarget. 2017;8(42):72096–107.

70. Kurscheid S, Bady P, Sciuscio D, Samarzija I, Shay T, Vassallo I, et al. Chromosome 7 gain and DNA hypermethylation at the HOXA10 locus are associated with expression of a stem cell related HOX-signature in glioblastoma. Genome Biol. 2015;16:16.

71. Fukushima T, Takeshima H, Kataoka H. Anti-glioma therapy with temozolomide and status of the DNA-repair gene MGMT. Anticancer Res. 2009;29(11):4845–54.

72. Stupp R, Mason WP, van den Bent MJ, Weller M, Fisher B, Taphoorn MJB, et al. Radiotherapy plus Concomitant and Adjuvant Temozolomide for Glioblastoma. New England Journal of Medicine. 2005;352(10):987–96.

73. Li YH, Yu CY, Li XX, Zhang P, Tang J, Yang Q, et al. Therapeutic target database update 2018: enriched resource for facilitating bench-to-clinic research of targeted therapeutics. Nucleic Acids Res. 2018;46(D1):D1121–D7.

74. Noushmehr H, Weisenberger DJ, Diefes K, Phillips HS, Pujara K, Berman BP, et al. Identification of a CpG island methylator phenotype that defines a distinct subgroup of glioma. Cancer Cell. 2010;17(5):510–22.

75. Henrichsen CN, Chaignat E, Reymond A. Copy number variants, diseases and gene expression. Hum Mol Genet. 2009;18(R1):R1–8.

76. John CG Spainhour HSL, Soojin V Yi and Peng Qiu. Correlation Patterns Between DNA Methylation and Gene Expression in The Cancer Genome Atlas. Cancer Informatics. 2019;18:1176935119828776.

77. Han S, Xia J, Qin X, Han S, Wu A. Phosphorylated SATB1 is associated with the progression and prognosis of glioma. Cell Death Dis. 2013;4:e901.

78. Liu M, Xu Z, Du Z, Wu B, Jin T, Xu K, et al. The Identification of Key Genes and Pathways in Glioma by Bioinformatics Analysis. J Immunol Res. 2017;2017:1278081.

79. Bourgonje AM, Verrijp K, Schepens JTG, Navis AC, Piepers JAF, Palmen CBC, et al. Comprehensive protein tyrosine phosphatase mRNA profiling identifies new regulators in the progression of glioma. Acta Neuropathologica Communications. 2016;4(1).

80. Zhang M, Zhao Y, Zhao J, Huang T, Wu Y. Impact of AKAP6 polymorphisms on Glioma susceptibility and prognosis. BMC Neurology. 2019;19(1).

81. Wang L, He S, Tu Y, Ji P, Zong J, Zhang J, et al. Downregulation of chromatin remodeling factor CHD5 is associated with a poor prognosis in human glioma. J Clin Neurosci. 2013;20(7):958–63.

82. Kennedy NJ, Cellurale C, Davis RJ. A radical role for p38 MAPK in tumor initiation. Cancer Cell. 2007;11(2):101–3.

83. Wong JC, Fiscus RR. Essential roles of the nitric oxide (no)/cGMP/protein kinase G type-Iα (PKG-Iα) signaling pathway and the atrial natriuretic peptide (ANP)/cGMP/PKG-Iα autocrine loop in promoting proliferation and cell survival of OP9 bone marrow stromal cells. Journal of Cellular Biochemistry. 2011;112(3):829–39.

84. Gravendeel LA, Kouwenhoven MC, Gevaert O, de Rooi JJ, Stubbs AP, Duijm JE, et al. Intrinsic gene expression profiles of gliomas are a better predictor of survival than histology. Cancer Res. 2009;69(23):9065–72.

85. Sturm D, Witt H, Hovestadt V, Khuong-Quang DA, Jones DT, Konermann C, et al. Hotspot mutations in H3F3A and IDH1 define distinct epigenetic and biological subgroups of glioblastoma. Cancer Cell. 2012;22(4):425–37.

86. Sun L, Hui AM, Su Q, Vortmeyer A, Kotliarov Y, Pastorino S, et al. Neuronal and glioma-derived stem cell factor induces angiogenesis within the brain. Cancer Cell. 2006;9(4):287–300.

87. Griesinger AM, Birks DK, Donson AM, Amani V, Hoffman LM, Waziri A, et al. Characterization of distinct immunophenotypes across pediatric brain tumor types. J Immunol. 2013;191(9):4880–8.

88. Gill BJ, Pisapia DJ, Malone HR, Goldstein H, Lei L, Sonabend A, et al. MRI-localized biopsies reveal subtype-specific differences in molecular and cellular composition at the margins of glioblastoma. Proc Natl Acad Sci U S A. 2014;111(34):12550–5.

89. Bredel M, Bredel C, Juric D, Harsh GR, Vogel H, Recht LD, et al. Functional network analysis reveals extended gliomagenesis pathway maps and three novel MYC-interacting genes in human gliomas. Cancer Res. 2005;65(19):8679–89.

90. Bredel M, Bredel C, Juric D, Duran GE, Yu RX, Harsh GR, et al. Tumor necrosis factor-alpha-induced protein 3 as a putative regulator of nuclear factor-kappaB-mediated resistance to O6-alkylating agents in human glioblastomas. J Clin Oncol. 2006;24(2):274–87.

91. Yan W, Zhang W, You G, Zhang J, Han L, Bao Z, et al. Molecular classification of gliomas based on whole genome gene expression: a systematic report of 225 samples from the Chinese Glioma Cooperative Group. Neuro-oncology. 2012;14(12):1432–40.

92. Lai RK, Chen Y, Guan X, Nousome D, Sharma C, Canoll P, et al. Genome-wide methylation analyses in glioblastoma multiforme. PLoS One. 2014;9(2):e89376.

93. Mur P, Mollejo M, Ruano Y, de Lope AR, Fiano C, Garcia JF, et al. Codeletion of 1p and 19q determines distinct gene methylation and expression profiles in IDH-mutated oligodendroglial tumors. Acta Neuropathol. 2013;126(2):277–89.

94. Zhang W, Yan W, You G, Bao Z, Wang Y, Liu Y, et al. Genome-wide DNA methylation profiling identifies ALDH1A3 promoter methylation as a prognostic predictor in G-CIMP-primary glioblastoma. Cancer Lett. 2013;328(1):120–5.

